# Enteroendocrine cells sense bacterial tryptophan catabolites to activate enteric and vagal neuronal pathways

**DOI:** 10.1101/2020.06.09.142133

**Authors:** Lihua Ye, Munhyung Bae, Chelsi D. Cassilly, Sairam V. Jabba, Daniel W. Thorpe, Alyce M Martin, Hsiu-Yi Lu, Jinhu Wang, John D. Thompson, Colin R. Lickwar, Kenneth D. Poss, Damien J. Keating, Sven-Eric Jordt, Jon Clardy, Rodger A. Liddle, John F. Rawls

## Abstract

The intestinal epithelium senses nutritional and microbial stimuli using epithelial sensory enteroendocrine cells (EECs). EECs can communicate nutritional information to the nervous system, but similar mechanisms for microbial information are unknown. Using *in vivo* real-time measurements of EEC and nervous system activity in zebrafish, we discovered that the bacteria *Edwardsiella tarda* specifically activates EECs through the receptor transient receptor potential ankyrin A1 (Trpa1) and increases intestinal motility in an EEC-dependent manner. Microbial, pharmacological, or optogenetic activation of Trpa1^+^EECs directly stimulates vagal sensory ganglia and activates cholinergic enteric neurons through 5-HT. We identified a subset of indole derivatives of tryptophan catabolism produced by *E. tarda* and other gut microbes that potently activates zebrafish EEC Trpa1 signaling and also directly stimulates human and mouse Trpa1 and intestinal 5-HT secretion. These results establish a molecular pathway by which EECs regulate enteric and vagal neuronal pathways in response to specific microbial signals.

## INTRODUCTION

The intestine harbors complex microbial communities that shape intestinal physiology, modulate systemic metabolism, and regulate brain function. These effects on host biology are often evoked by distinct microbial stimuli including microbe-associated molecular patterns (MAMPs) and microbial metabolites derived from digested carbohydrates, proteins, lipids, and bile acids (Brown and Hazen, 2015, Liu et al., 2020, Medzhitov, 2007). The intestinal epithelium is the primary interface that mediates this host-microbe communication (Kaiko and Stappenbeck, 2014). The mechanisms by which the intestinal epithelium senses distinct microbial stimuli and transmits that information to the rest of the body remains incompletely understood.

The intestinal epithelium has evolved specialized enteroendocrine cells (EECs) that exhibit conserved sensory functions in insects, fishes, and mammals (Guo et al., 2019, Ye et al., 2019, Furness et al., 2013). Distributed along the entire digestive tract, EECs are activated by diverse luminal stimuli to secrete hormones or neuronal transmitters in a calcium dependent manner (Furness et al., 2013). Recent studies have revealed that EECs form synaptic connections with sensory neurons (Kaelberer et al., 2018, Bellono et al., 2017, Bohorquez et al., 2015). The connection between EECs and neurons forms a direct route for the intestinal epithelium to transmit nutrient sensory information to the brain (Kaelberer et al., 2018). EECs are classically known for their ability to sense nutrients (Symonds et al., 2015) but whether they can be directly stimulated by microbes or microbially derived products is less clear. Limited examples include the observation that short chain fatty acids and branched chain fatty acids from microbial carbohydrate and amino acid catabolism activate EECs via G-protein coupled receptors (Bellono et al., 2017, Lu et al., 2018). Indole, a microbial catabolite of the amino acid tryptophan, has also been reported to activate EECs, but the EEC receptor that mediates the effect remains unidentified (Chimerel et al., 2014). With the growing understanding of gut microbiota and their metabolites, identifying the EEC receptors that recognize distinct microbial stimuli as well as the downstream pathways by which EECs transmit microbial stimuli to regulate local and systemic host physiology, has emerged as an important goal.

The vertebrate intestine is innervated by the intrinsic enteric nervous system (ENS) and extrinsic neurons from autonomic nerves, including sensory nerve fibers from the nodose vagal ganglia and dorsal root ganglia in the spinal cord (Furness et al., 1999). Both vagal and spinal sensory nerve fibers transmit visceral stimuli to the central nervous system and modulate a broad spectrum of brain functions (Brookes et al., 2013). A previous study demonstrated that stimulating EECs with the microbial metabolite isovalerate activates spinal sensory nerves through 5-hydroxytryptamine (5-HT) secretion (Bellono et al., 2017). Whether and how gut microbial stimuli modulate ENS or vagal sensory activity through EECs is still unknown.

EECs are known to express a broad diversity of receptors and channels to perceive and respond to environmental stimuli (Furness et al., 2013). Transient receptor potential ankyrin 1 (Trpa1) is an excitatory calcium-permeable non-selective cation channel that can be activated by multiple chemical irritants and has important roles in pain sensation and neurologic inflammation (Bautista et al., 2013, Lapointe and Altier, 2011). Many of the known Trpa1 agonists are chemicals derived from food spices or environmental pollution (Nilius et al., 2011). Whether microbial metabolites also activate Trpa1 is completely unknown.

Here, we show that Trpa1 is expressed in a subset of EECs that can be uniquely activated by gut microbes. Specifically, we identified a gram-negative bacterium, *Edwardsiella tarda* (*E. tarda*), that activates EECs in a Trpa1 dependent manner. Microbial, optochemical, or optogenetic activation of Trpa1+EECs activates vagal sensory ganglia and increases intestinal motility through direct signaling to enteric motor neurons. We also identified a subset of tryptophan catabolites that secreted from *E. tarda* and other gut microbes that potently activate Trpa1, stimulate intestinal motility, and activate vagal neurons. These results establish a molecular pathway by which EECs regulate enteric and vagal neuronal activity in response to specific microbial signals in the gut.

## RESULTS

### *Edwardsiella tarda* activates EECs through Trpa1

To identify stimuli that activate EECs in live animals, we developed a new transgenic zebrafish line that permits recording of EEC activity by expressing the calcium modulated photoactivatable ratiometric integrator (CaMPARI) protein in EECs under control of the *neurod1* promoter (Fig. 1A, Fig. S1A-F; see Methods for details). When exposed to 405nm light, CaMPARI protein irreversibly photoconverts from a configuration that emits green light to one that emits red in a manner positively correlated with intracellular calcium levels [Ca^2+^]_i_. A high red:green CaMPARI ratio thus reports high intracellular calcium (Fosque et al., 2015). This EEC-CaMPARI system therefore enables imaging of the calcium activity history of intestinal EECs in the intact physiologic context of live free-swimming animals (Fig. 1A-B, Fig. S1G-J). To test the validity of this EEC-CaMPARI system, we first stimulated larvae with different nutrients known to activate zebrafish EECs (Ye et al., 2019). Exposure to only water as a vehicle control revealed an expected low basal red:green CaMPARI ratio (Fig. 1C, E-F). Following long-chain fatty acid stimulation with linoleate, a subpopulation of EECs displayed high red:green CaMPARI ratio (Fig. 1D, E-F). EECs with a high red:green CaMPARI ratio were classified as “activated EECs”. The percentage of activated EECs significantly increased in response to chemical stimuli known to activate EECs, including linoleate, oleate, laurate, and glucose (Fig. 1F), but not in response to the short chain fatty acid butyrate, consistent with our previous finding (Fig. 1F) (Ye et al., 2019).

**Figure 1.**
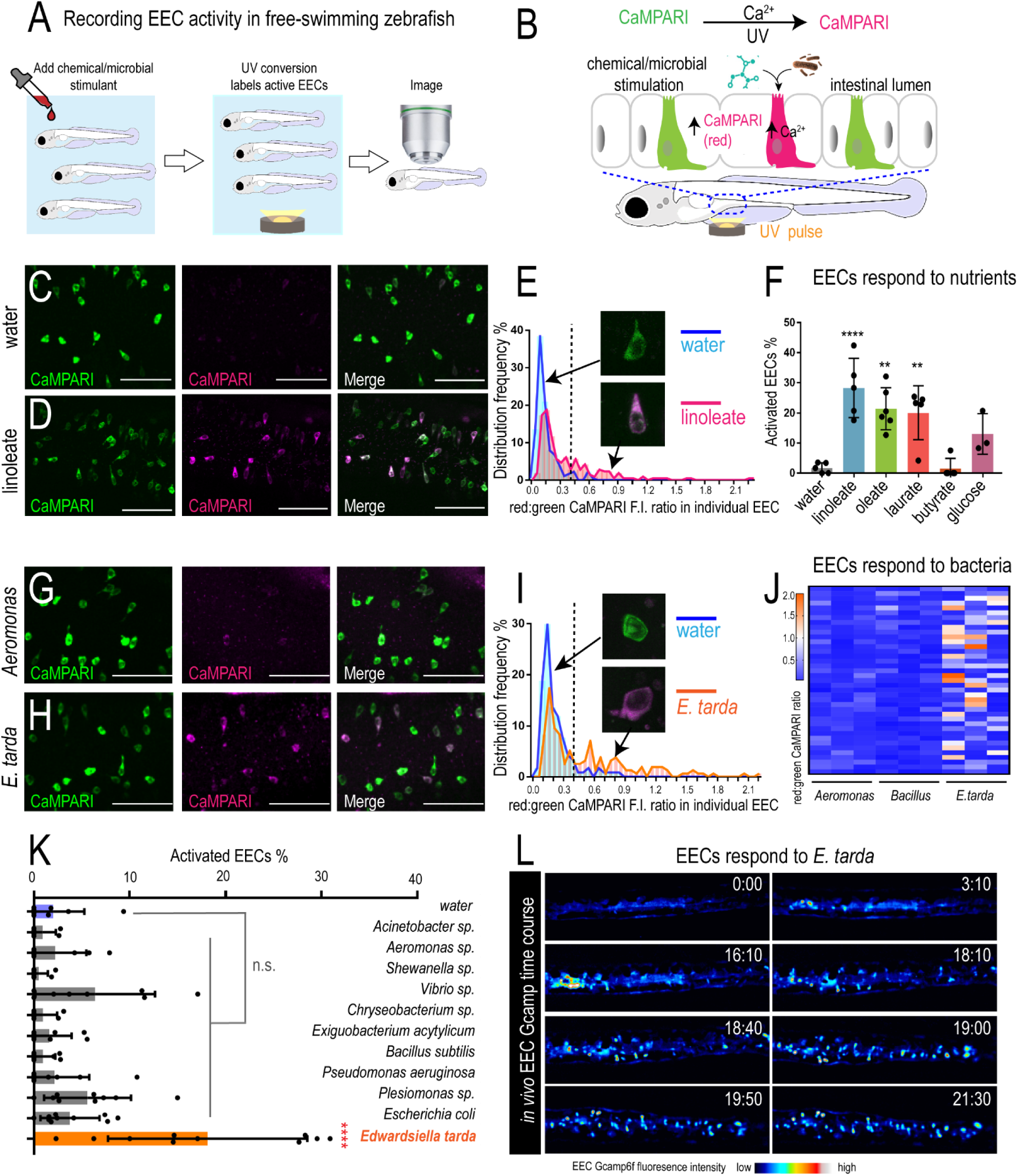
*E. tarda* activates zebrafish EECs *in vivo*. (A) Experimental approach for measuring EEC activity in free-swimming zebrafish. (B) Method for recording EEC responses to chemical and microbial stimulants in the EEC-CaMPARI model. (C-D) Confocal projection of mid-intestinal EECs upon water (C, negative control) or linoleate (D) stimulation in *Tg(neurod1:CaMPARI)* following UV-photoconversion. (E) Frequency distribution of EECs’ red:green CaMPARI fluorescence intensity ratio in water or linoleate-stimulated zebrafish. n=177 for water group and n=213 for linoleate group. (F) Percent EEC response in *Tg(neurod1:CaMPARI)* zebrafish. (G-H) Confocal projection of mid-intestinal EECs upon *Aeromonas* sp. (G) or *E. tarda* (H) stimulation in *Tg(neurod1:CaMPARI)* following UV-photoconversion. (I) Frequency distribution of EECs’ red:green CaMPARI fluorescence intensity ratio in zebrafish treated with water or *E. tarda*. n=117 for water group and n=156 for *E. tarda* group. (J) Representative heatmap image showing *Aeromonas* sp., *B. subtilis* and *E. tarda* stimulated EEC red:green CaMPARI fluorescence ratio. (K) EEC activation in *Tg(neurod1:CaMPARI)* zebrafish stimulated with different bacterial strains. (L) Representative *Tg(neurod1:Gcamp6f)* zebrafish intestine stimulated with *E. tarda*. One-way ANOVA with Tukey’s post-test was used in F and K. *p<0.05; **p<0.01; ***p<0.001; ****p<0.0001.

We next applied the EEC-CaMPARI system to ask whether EECs acutely respond to live bacterial stimulation *in vivo*. We exposed *Tg(neurod1:CaMPARI)* zebrafish to individual bacterial strains for 20 mins in zebrafish housing water (GZM), followed by photoconversion and imaging of CaMPARI fluorescence. For these experiments, we selected a panel of 11 bacterial strains including 3 model species (*P. aeruginosa, E. coli, B. subtilis*), 7 commensal strains isolated from the zebrafish intestine (Rawls et al., 2006, Stephens et al., 2016), and the pathogen *E. tarda* FL6-60 (also called *E. piscicida* (Abayneh et al., 2013, Bujan et al., 2018); Fig. 1K and Key Resources Table). Within this panel, the only strain that induced a high red:green EEC-CaMPARI signal was *E. tarda* (Fig. 1G-K). We further confirmed that *E. tarda* directly activates EECs using an alternative reporter of EEC activity based on the [Ca^2+^]_i_-sensitive fluorescent protein Gcamp6f (*neurod1:Gcamp6f*) (Fig. 1L, Fig. S1K-P and Video 1) (Ye et al., 2019). Although *E. tarda* has been reported to infect zebrafish (Abayneh et al., 2013, Flores et al., 2020), we observed no overt pathogenesis in these acute exposure experiments.

EECs express a variety of sensory receptors that can be activated by different environmental stimuli. To investigate the molecular mechanisms by which EECs perceive *E. tarda* stimulation, we isolated EECs from zebrafish larvae and performed RNA-seq analysis. Transcript levels in FACS-sorted EECs (*cldn15la:EGFP*+; *neurod1:TagRFP*+) were compared to all other intestinal epithelial cells (IECs) (*cldn15la:EGFP*+; *neurod1:TagRFP*-) (Fig. 2A). We identified 192 zebrafish transcripts that were significantly enriched in EECs by DESeq2 using P_FDR_<0.05 (Fig. 2B and Table S1). Gene Ontology (GO) term analysis revealed that those zebrafish genes are enriched for processes like hormone secretion, chemical synaptic transmission and neuropeptide signaling (Table S1). To identify gene homologs enriched in EECs in both zebrafish and mammals, we compared these 192 genes to published RNA-seq data from *Neurod1*+ EECs from mouse duodenum and CHGA+ EECs from human jejunum (Roberts et al., 2019). Despite the evolutionary distance and differences in tissue origin, we found that 24% of zebrafish EEC-enriched gene homologs (46 out of 192) were shared among zebrafish, human, and mouse, and that 40% of zebrafish EEC-enriched genes (78 out of 192) were shared between zebrafish EECs and human jejunal EECs (Table S2). The genes with conserved EEC expression include those encoding specific hormones, transcription factors, G-protein coupled receptors, and ion channels that regulate membrane potential (Fig. 2C and Table S3). Using published data from mouse intestinal epithelial single-cell RNA-seq data that revealed different EEC subtypes (Haber et al., 2017), we found that many of the signature genes in mouse enterochromaffin cells (EC), which are identified by their 5-HT synthesis, are enriched in zebrafish EECs (Table S3). Among these conserved EEC-enriched genes, one of the genes with the highest expression in zebrafish EECs is transient receptor potential ankyrin 1 (Trpa1) (Fig. 2C and Table S1, 2).

**Figure 2.**
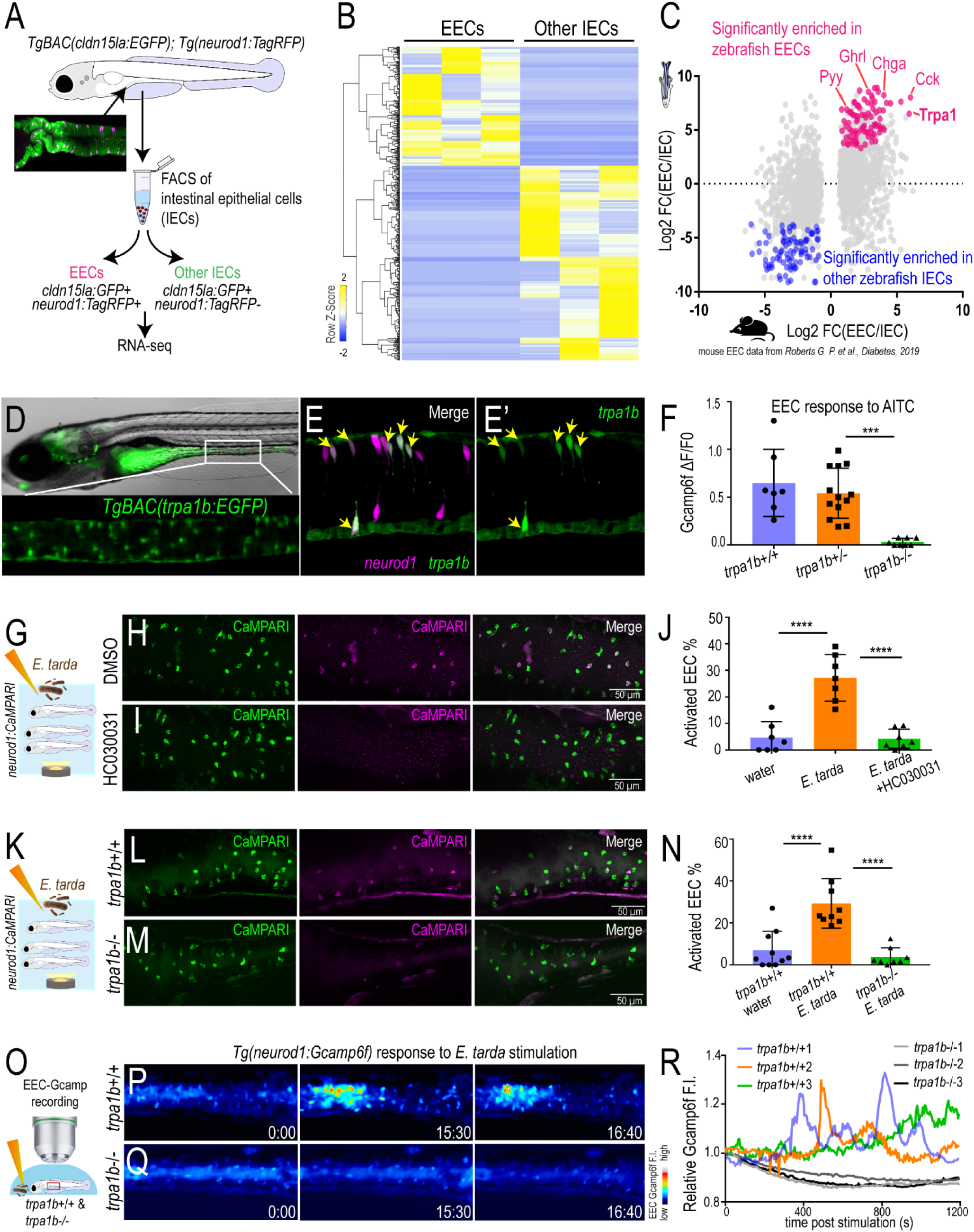
*E. tarda* activates EECs through Trpa1. (A) Schematic diagram of zebrafish EEC RNA-seq. (B) Clustering of genes that are significantly enriched in zebrafish EECs and other IECs (P_adj_<0.05). (C) Comparison of zebrafish and mouse EEC enriched genes. Mouse EEC RNA-seq data was obtained from GSE114913 (Roberts et al., 2019). (D) Fluorescence image of *TgBAC(trpa1b:EGFP)*. Zoom-in view shows the expression of *trpa1b+* cells in intestine. (E) Confocal projection of a *TgBAC(trpa1b:EGFP);Tg(neurod1:TagRFP)* zebrafish intestine. Yellow arrows indicate zebrafish EECs that are *trpa1b:EGFP*+. (F) Quantification of EEC Gcamp responses to Trpa1 agonist AITC stimulation in *trpa1b*+/+, *trpa1b*+/- and *trpa1b*-/-zebrafish. (G) Experimental design. (H-I) Confocal projection of *Tg(neurod1:CaMPARI)* zebrafish intestine stimulated with *E. tarda* with or without the Trpa1 antagonist HC030031. (J) Quantification of activated EECs in control and HC030031 treated zebrafish treated with water or *E. tarda*. (K) Experimental approach. (L-M) Confocal projection of *trpa1b*+/+ or *trpa1b*-/-*Tg(neurod1:CaMPARI)* intestine after stimulation with water or *E. tarda*. (N) Quantification of activated EEC percentage in WT and *trpa1b*-/-zebrafish treated with water or *E. tarda*. (O) Experimental design. (P-Q) Timed images of *trpa1b*+/+ or *trpa1b*-/-*Tg(neurod1:Gcamp6f)* zebrafish stimulated with *E. tarda*. (R) Quantification of relative EEC Gcamp6f fluorescence intensity in WT or *trpa1b*-/-zebrafish treated with *E. tarda*. One-way ANOVA with Tukey’s post-test was used in F, J, N. *p<0.05; **p<0.01; ***p<0.001; ****p<0.0001.

The zebrafish genome encodes two *trpa1* paralogs, *trpa1a* and *trpa1b* (Prober et al., 2008). Zebrafish EECs express *trpa1b* but not *trpa1a*, as revealed by our RNA-seq data (Fig. S2A) and RT-PCR confirmation in FACS-isolated EECs (Fig. S2B). Fluorescence imaging of *TgBAC(trpa1b:EGFP)* zebrafish (Pan et al., 2012) further revealed that *trpa1b* is expressed within the intestinal epithelium by a distinct subset of cells expressing the EEC marker *neurod1* (Fig. 2D-E). In addition, zebrafish EECs were activated by exposure to the Trpa1 agonist allyl isothiocyanate (AITC) (Fig. 2F, Fig. S2C, G and Video 2), whereas this response was inhibited by the Trpa1 antagonist HC030031 (Fig. S2D). AITC was unable to induce EEC activation in zebrafish homozygous for a *trpa1b* mutation but was able to induce EEC activation normally in *trpa1a* mutants (Prober et al., 2008) (Fig. 2F, Fig. S2E-F and Video 2). These data establish that *trpa1b*, but not *trpa1a*, is expressed by a subset of zebrafish EECs and is required for EEC activation by Trpa1 agonist AITC.

Trpa1 is a nociception receptor that is known to mediate pain sensation in nociceptive neurons (Lapointe and Altier, 2011). A broad spectrum of chemical irritants, including many compounds that are derived from food spices, activate Trpa1 (Nilius et al., 2011). In addition to chemical irritants, certain bacterial products, including lipopolysaccharide (LPS) and hydrogen sulfide (H_2_S), stimulated nociceptive neurons in a Trpa1-dependent manner (Meseguer et al., 2014). Since the expression of classic microbial pattern recognition receptors is very low in zebrafish EECs (Table S3), we tested if Trpa1 mediated *E. tarda*-induced EEC activation. We first observed that treatment of wild-type (WT) *Tg(neurod1:CaMPARI)* fish with the Trpa1 antagonist HC030031 significantly inhibited *E. tarda*’s ability to induce EEC activation (Fig. 2G-J). The ability of *E. tarda* to induce EEC activity in the EEC-CaMPARI model was similarly blocked in *trpa1b* mutant zebrafish (Fig. 2K-N). In accord, experiments in *Tg(neurod1:Gcamp6f)* zebrafish confirmed that *Gcamp6f* fluorescence increased in EECs in response to *E. tarda* stimulation in WT, but not *trpa1b* mutant zebrafish (Fig. 2O-R). Therefore live *E. tarda* bacteria stimulate EECs in a Trpa1-dependent manner, suggesting that EEC Trpa1 signaling may play an important role in mediating microbe-host interactions.

### EEC Trpa1 signaling is important to maintain microbial homeostasis by regulating intestinal motility

To determine how *E. tarda*-induced Trpa1 signaling in EECs affects the host, we exposed *trpa1b*^*+/+*^ and *trpa1b*^-/-^ zebrafish larvae to an *E. tarda* strain that expresses mCherry fluorescent protein. High-dose (10^7^ CFU/mL) *E. tarda* exposure for 3 days decreased survival rate and caused gross pathology (Fig. S2M-N), consistent with its reported activity as a zebrafish pathogen (Abayneh et al., 2013, Flores et al., 2020). To investigate the interaction between *E. tarda* and Trpa1+EECs under relatively normal physiological conditions, we exposed zebrafish with a low *E. tarda* dose (10^6^ CFU/mL) that did not significantly affect survival rate or cause gross pathology (Fig. S2M-N and Fig. 3A). When reared under conventional conditions in the absence of *E. tarda*, we observed no significant difference in the abundance of culturable gut microbes between *trpa1b*^+/+^ and *trpa1b*^-/-^ zebrafish (Fig. S2H-I). However, upon 3-day treatment with *E. tarda*, there was significant accumulation of *E. tarda* mCherry^+^ bacteria in the intestinal lumen in *trpa1b*^-/-^ but not *trpa1b*^+/+^ zebrafish larvae (Fig. 3A-C). This accumulation could be observed by either quantifying *E. tarda* mCherry fluorescence (Fig. 3D) or counting *E. tarda* colony forming units (CFU) from digestive tracts dissected from *E. tarda* treated *trpa1b*^+/+^ and *trpa1b*^-/-^ zebrafish (Fig. 3E). This suggests that Trpa1 signaling may act as a host defense mechanism to facilitate clearance of specific types of bacteria such as *E. tarda*.

**Figure 3.**
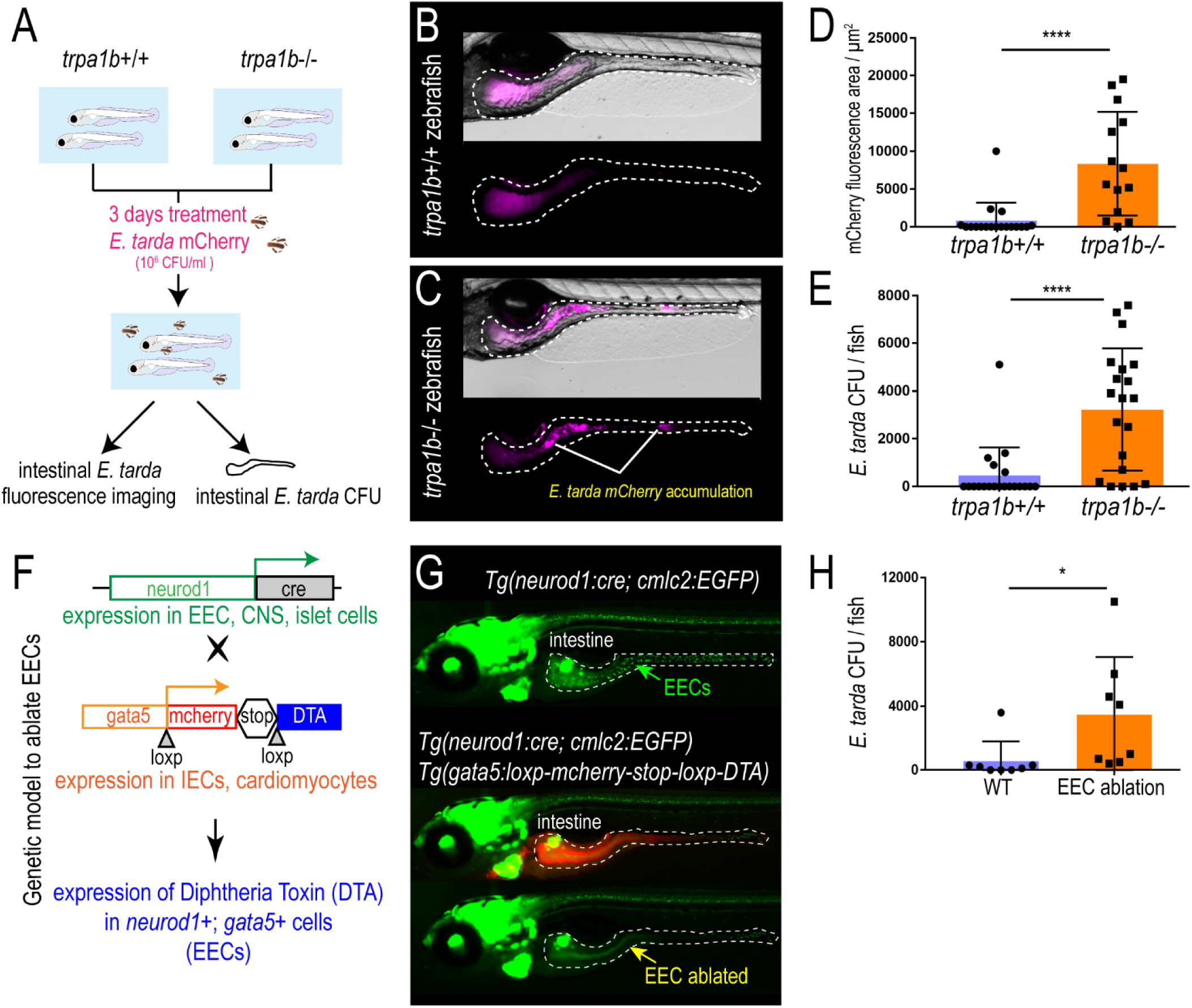
Activation of EEC Trpa1 signaling facilitates enteric *E. tarda* clearance. (A) Schematic of zebrafish *E. tarda* treatment. (B-C) Representative image of *trpa1b*+/+ or *trpa1b*-/-zebrafish treated with *E. tarda* expressing mCherry (*E. tarda* mCherry). (D) Quantification of *E. tarda* mCherry fluorescence in *trpa1b*+/+ or *trpa1b*-/-zebrafish intestine. (E) Quantification of intestinal *E. tarda* CFU in *trpa1b*+/+ or *trpa1b*-/-zebrafish. (F) Schematic of genetic model in which EECs are ablated via Cre-induced Diptheria Toxin (DTA) expression. (G) Representative image of *Tg(neurod1:cre; cmlc2:EGFP)* and *Tg(neurod1:cre; cmlc2:EGFP); TgBAC(gata5:RSD)* with EECs that are labelled by *Tg(neurod1:EGFP)*. (H) Quantification of intestinal *E. tarda* CFU in WT or EEC ablated zebrafish. Student’s t-test was used in D, E, H. *p<0.05; ****p<0.0001.

In addition to EECs, Trpa1 is also expressed in mesenchymal cells within the intestine (Fig.2D-E and Fig. S4O) and nociceptive sensory neurons (Yang et al., 2019, Holzer, 2011). To investigate whether the phenotype we observed above is specifically mediated by EECs, we generated a new Cre-*loxP* transgenic system that permits specific ablation of EECs without affecting other IECs or other *neurod1* expressing cells like CNS or pancreatic islets (Fig. 3F, Fig. S2J). Quantitative RT-PCR and immunofluorescence confirmed a reduction of EEC hormones but not non-EEC marker genes (Fig. S2K-L). Establishing this EEC ablation system allowed us to define the specific role of EECs in mediating *E. tarda*-host interaction. As with *trpa1b*^-/-^ zebrafish, we did not detect significant differences in gut microbial abundance between unexposed WT and EEC-ablated zebrafish (Fig. S2O). However, in response to *E. tarda* exposure, a significantly higher amount of *E. tarda* mCherry accumulated in EEC-ablated zebrafish compared to WT sibling controls (Fig. 3H and Fig. S2P-Q). Together, these data establish that EEC Trpa1 signaling maintains gut microbial homeostasis by facilitating host clearance of specific types of bacteria like *E. tarda*.

To understand the mechanisms by which EEC Trpa1 regulates gut microbial homeostasis, we used an opto-pharmacological approach that permits temporal control of EEC Trpa1 activation through UV light exposure (Fig. 4A). We pretreated zebrafish with Optovin, a chemical that specifically activates Trpa1 only in the presence of UV light (Kokel et al., 2013) (Fig. S3A). To specifically activate Trpa1 in EECs, we mounted zebrafish larvae pretreated with Optovin and restricted UV light exposure specifically to the intestinal epithelium using a confocal laser (Fig. S3A). UV light activation significantly increased [Ca^2+^]_i_ in a subpopulation of EECs in WT larvae, as measured by Gcamp6f fluorescence (Fig. S3B-C, Video 3). The same UV light exposure in *trpa1b*^-/-^ larvae pretreated with Optovin did not increase EEC [Ca^2+^]_i_ (Fig. 4B-C), indicating that EEC activation induced by Optovin-UV was dependent on Trpa1. Next, we used this approach to examine the effect of EEC Trpa1 activation on intestinal motility. Trpa1 activation in EECs within the middle intestine via UV light application in WT larvae produced a propulsive movement of the intestine from anterior to posterior, and the velocity of intestinal motility increased accordingly (Fig. 4D-F, Fig. S3D-E and Video 4). In contrast, Optovin treatment and UV activation failed to induce intestinal motility in EEC-ablated zebrafish (Fig. 4D-F and Video 5). These results indicate that intestinal motility triggered by Trpa1 activation is dependent on EECs. To further test if signaling from Trpa1+EECs is sufficient to activate intestinal motility, we developed a new optogenetic system in which a mCherry tagged Channelrhodopsin (ChR2-mCherry) is expressed in EECs from the *neurod1* promoter (Fig. 4G-H). Blue light activation of ChR2 causes cation influx and plasma membrane depolarization, and [Ca^2+^]_i_ then increases through the activation of voltage-dependent calcium channels (Nagel et al., 2003) which are abundantly expressed in zebrafish EECs (Fig. 4I-J, Table S3 and Video 6). This new tool permits selective activation of the ChR2-mCherry+ EECs using a confocal laser, without affecting the activity of nearby EECs (see Methods and Fig. S3F). We therefore used *Tg(neurod1:Gal4); Tg(UAS:ChR2-mCherry); TgBAC(trpa1b:EGFP)* larvae to selectively activate ChR2-mCherry expressing EECs that are either *trpa1b*+ or *trpa1b*-. We found that activation of *trpa1b*+ EECs but not *trpa1b*-EECs consistently increased intestinal velocity magnitude (Fig. 4K-L, Fig. S3F-H and Video 7, 8), again indicating a unique role for Trpa1+EECs in regulating intestinal motility. Consistent with the Optovin-UV result, stimulating Trpa1+ChR2+ EECs in the middle intestine resulted in anterograde intestinal movement (Video 8). Interestingly, stimulating Trpa1+ChR2+ EECs in the proximal intestine initiated a retrograde intestinal movement (Video 7). This is consistent with previous findings that the zebrafish proximal intestine typically exhibits a retrograde motility pattern whereas the middle and distal intestine display antegrade motility (Fig. S3D) (Roach et al., 2013). Finally, we tested whether microbial activation of Trpa1 signaling in EECs also increased intestinal motility. Using microgavage (Cocchiaro and Rawls, 2013), we found that delivery of live *E. tarda* into the intestinal lumen significantly promoted intestinal peristalsis and motility compared to PBS-gavaged controls (Fig. 4M-O and Video 9). *E. tarda* induced intestinal motility was significantly reduced in *trpa1b*-/-zebrafish (Fig. S3I). When we gavaged zebrafish with *Aeromonas* or *Bacillus*, two of the tested bacterial strains that do not activate EECs (Fig. 1), no significant change of intestinal motility was observed (Fig. 4M-O and Video 9). These experiments together establish that activation of Trpa1 in EECs directly stimulates intestinal motility, and provide a potential physiologic mechanism underlying Trpa1-dependent clearance of *E. tarda* from the intestinal lumen.

**Figure 4.**
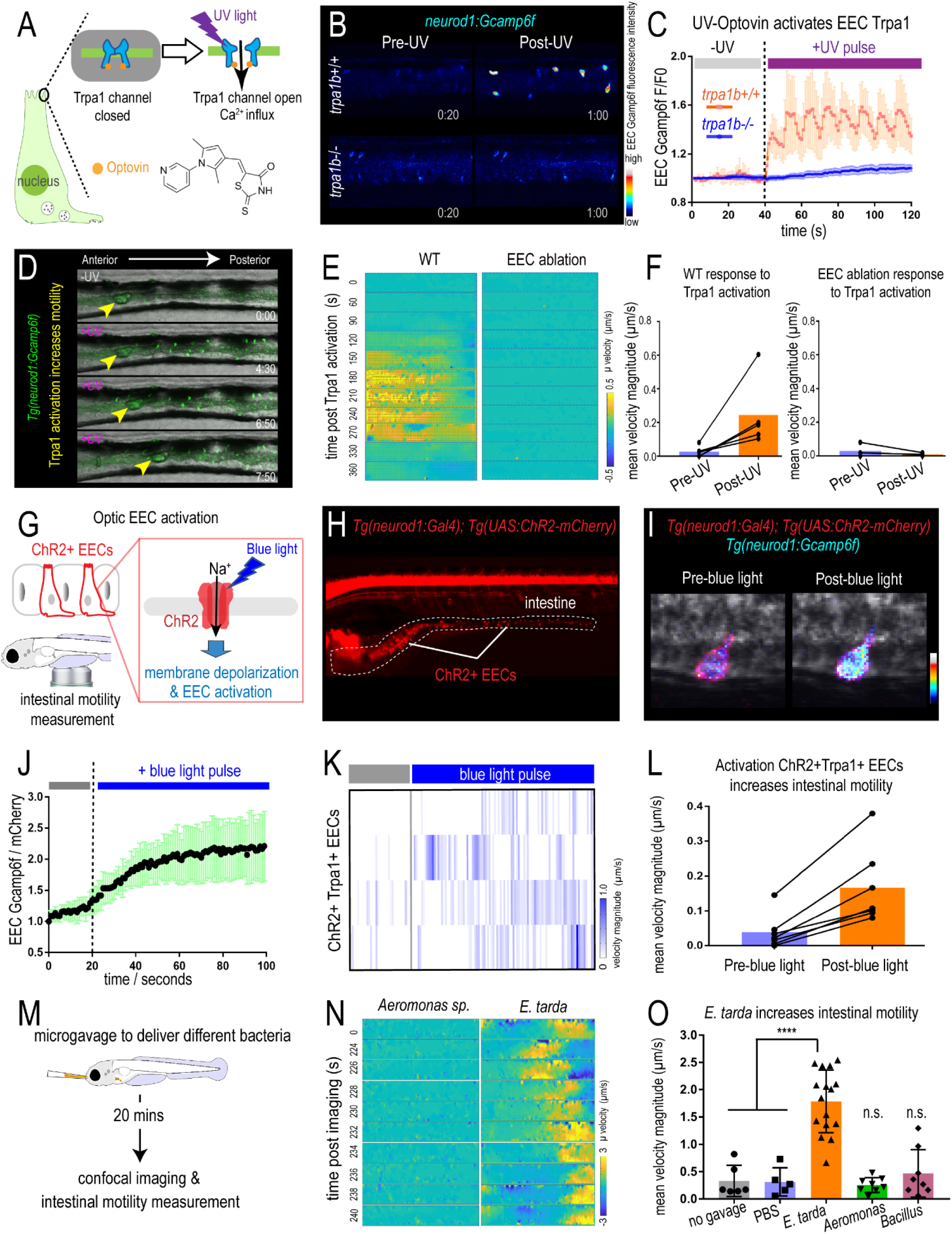
Activation of EEC Trpa1 signaling promotes intestinal motility. (A) Illustration of EEC Trpa1 activation using an Optovin-UV platform. (B) Confocal image of *trpa1b*+/+ and *trpa1b*-/-*Tg(neurod1:Gcamp6f)* zebrafish EECs before and after UV activation. (C) Quantification of EEC Gcamp6f fluorescence changes in *trpa1b*+/+ and *trpa1b*-/-zebrafish before and after UV induction. (D) Representative images of *Tg(neurod1:Gcamp6f)* zebrafish intestine before and after UV-induced Trpa1 activation. Yellow arrowheads indicate the movement of intestinal luminal contents from anterior to posterior following EEC activation. (E) PIV-Lab velocity analysis to quantify intestinal motility in WT and EEC ablated zebrafish. Spatiotemporal heatmap series represent the µ velocity of the imaged intestinal segment at the indicated timepoint post Trpa1 activation. (F) Quantification of the mean intestinal velocity magnitude before and after UV activation in WT and EEC ablated zebrafish. (G) Model of light activation of ChR2 in EECs. (H) Fluorescence image of *Tg(neurod1:Gal4); Tg(UAS:ChR2-mCherry)* zebrafish that express ChR2 in EECs. (I) Confocal image of ChR2 expressing EECs in *Tg(neurod1:Gcamp6f)* intestine before and after blue light-induced ChR2 activation. (J) Quantification of EEC Gcamp fluorescence intensity before and after blue light-induced ChR2 activation. (K) Intestinal velocity magnitude before and after blue-light induced activation in ChR2+Trpa1+ EECs. (L) Mean velocity magnitude before and after blue light-induced activation in ChR2+Trpa1+ EECs. (M) Experimental design schematic for panels N and O. (N) Heatmap representing the µ velocity of the imaged intestinal segment at indicated timepoints following *Aeromonas* sp. or *E. tarda* gavage. (O) Mean intestinal velocity magnitude in zebrafish without gavage or gavaged with PBS or different bacterial strains. Student’s t-test was used in O. ****p<0.0001.

### EEC Trpa1 signaling promotes intestinal motility by activating cholinergic enteric neurons

To test the role of the ENS in Trpa1-activated intestinal motility, we used zebrafish that lack an ENS due to mutation of the receptor tyrosine kinase gene *ret* (Taraviras et al., 1999). Immunofluorescence demonstrated that *ret*^-/-^ zebrafish lack all identifiable enteric nerves (marked by *NBT* transgenes, Fig. 5B and Fig. S4A-B), whereas EECs remain intact (marked by *neurod1* transgenes, Fig. 5B) and responsive to Trpa1 agonist (Fig. S4C-F). Using the Optovin-UV system (Fig. 5A), we observed that EEC Trpa1 activation increased intestinal motility in control (*ret*^+/+^ or *ret*^+/-^) but not *ret*^*-/-*^ zebrafish (Fig. 5C-D and Fig. S4G-H). These results were confirmed using a second zebrafish mutant that lacks an ENS due to mutation of the transcription factor gene *sox10* (Bondurand and Sham, 2013). Similar to *ret*^-/-^ zebrafish larvae, *sox10*^-/-^ zebrafish larvae lack an ENS but the EECs remain intact (Fig. S4I-L), and failed to increase intestinal motility following activation of EEC-Trpa1 signaling (Fig. S4M-N). These data suggest that Trpa1+ EECs do not signal directly to enteric smooth muscle to promote intestinal motility, but instead signal to the ENS.

**Figure 5.**
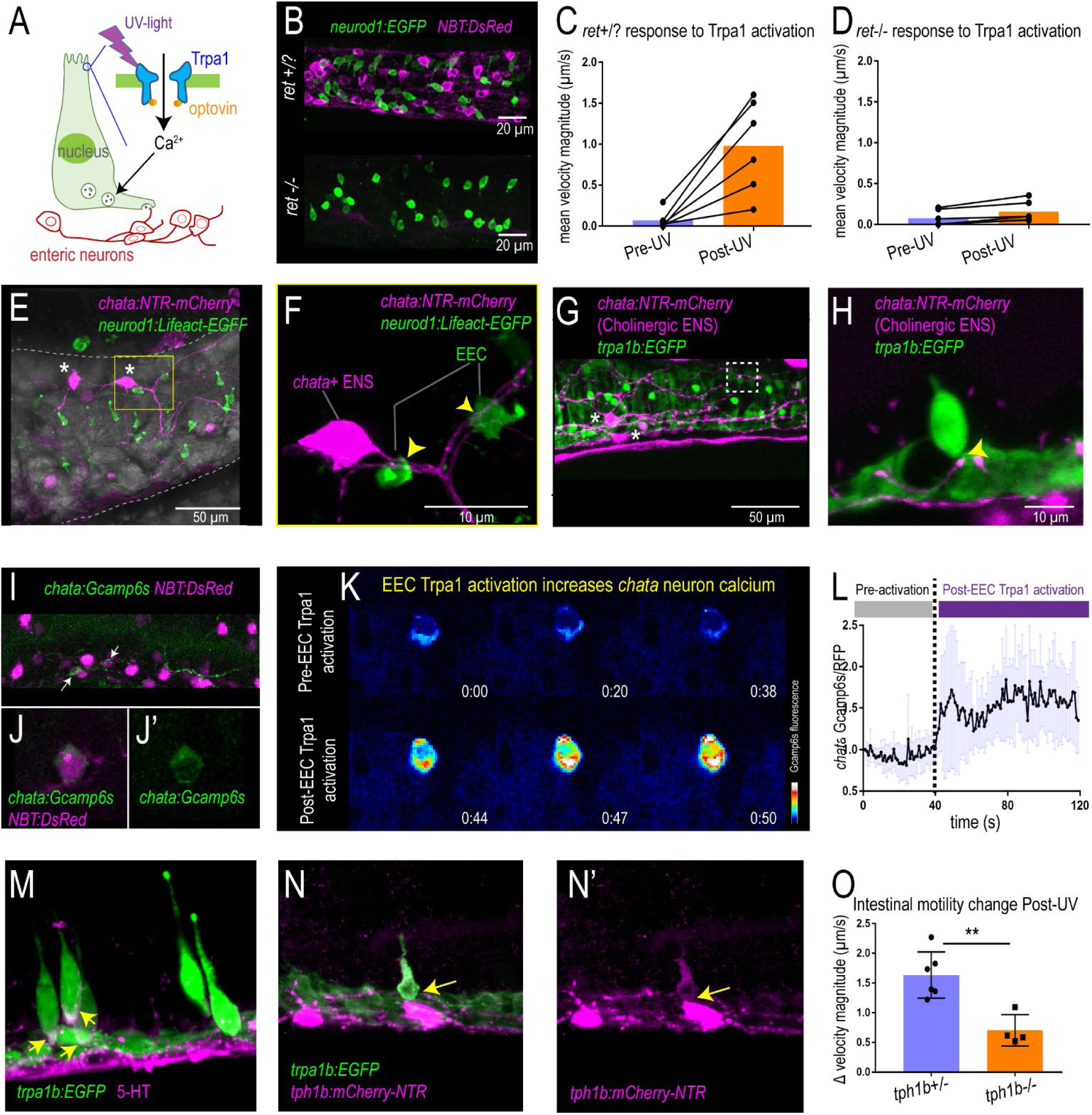
Activation of EEC Trpa1 signaling activates enteric cholinergic neurons and promotes intestinal motility through 5-HT. (A) Working model showing Trpa1 stimulation in EECs activates enteric neurons. (B) Confocal image of *ret*+/? (*ret*+/+ or *ret*+/-) and *ret*-/-intestine in *TgBAC(neurod1:EGFP);Tg(NBT:DsRed)* zebrafish. *neurod1* labelled EECs shown in green and *NBT* labelled ENS shown in magenta. (C) Quantification of mean intestinal velocity magnitude before and after EEC Trpa1 activation in *ret*+/? zebrafish. (D) Quantification of mean intestinal velocity magnitude before and after UV activation in *ret*-/-zebrafish. (E) Confocal image of intestine in *TgBAC(chata:Gal4); Tg(UAS:NTR-mCherry); Tg(neurod1:LifeAct-EGFP)* zebrafish. Cholinergic enteric neurons are shown in magenta and neurod1+ EECs are shown in green. The asterisks indicate Cholinergic enteric neuron cell bodies which reside on the intestinal wall. (F) Higher magnification view indicates the EECs (green) directly contact nerve fibers that are extended from the *chata*+ enteric neuron cell body (magenta). Yellow arrows indicate the points where EECs form direct connections with *chata*+ ENS. (G) Confocal image of intestine in *TgBAC(chata:Gal4); Tg(UAS:NTR-mCherry); TgBAC(trpa1b:EGFP)* zebrafish. (H) Trpa1+EECs (green) form direct contact with *chata*+ enteric neurons (magenta). (I) Live imaging of *TgBAC(chata:Gal4); Tg(UAS:Gcamp6s); Tg(NBT:DsRed)* zebrafish intestine. All the enteric neurons are labelled as magenta by *NBT:DsRed*. Yellow arrow indicates a *chata*+ enteric neuron. (J) Higher magnification view of a *Gcamp6s* expressing *chata*+ enteric neuron. (K) *In vivo* calcium imaging of *chata*+ enteric neuron before and after EEC Trpa1 activation. (L) Quantification of *chata*+ enteric motor neuron Gcamp6s fluorescence intensity before and after EEC Trpa1 activation. (M) Confocal image of *TgBAC(trpa1b:EGFP)* zebrafish intestine stained for 5-HT. Yellow arrows indicate the presence of 5-HT in the basal area of *trpa1b*+ EECs. (N) Confocal image of *TgBAC(trpa1b:EGFP);Tg(tph1b:mCherry-NTR)* zebrafish intestine. Yellow arrow indicates the *trpa1b*+ EECs that express Tph1b. (O) Quantification of intestinal motility changes in response to EEC Trpa1 activation in *tph1b*+/- and *tph1b*-/-zebrafish. Student’s t test was used in O. **p<0.01

The ENS is a complex network composed of many different neuronal subtypes. Among these subtypes, cholinergic neurons secrete the excitatory neurotransmitter acetylcholine to stimulate other enteric neurons or smooth muscle (Pan and Gershon, 2000, Qu et al., 2008). The cholinergic neurons are essential for normal intestinal motility (Johnson et al., 2018). One of the key enzymes for the synthesis of acetylcholine in the ENS is choline acetyltransferase (Chat) (Furness et al., 2014). Using *TgBAC(chata:Gal4); Tg(UAS:NTR-mCherry)* transgenic zebrafish, we were able to visualize the cholinergic enteric neurons in the zebrafish intestine (Fig. 5E and Fig. S5E-J). We found that *chata*+ neurons have smooth cell bodies which are located within the intestinal wall, many of which display multiple axons (Fig. 5E and Fig. S5E-F). Such multipolar neurons have also been classified as Dogiel type II neurons (Cornelissen et al., 2000). These Dogiel type II neurons are likely to be the intestinal intrinsic sensory neurons (Bornstein, 2006). We used *Tg(neurod1:LifeAct-EGFP); TgBAC(chata:Gal4); Tg(UAS:NTR-mCherry)* zebrafish to reveal that many EECs labeled by *neurod1* form direct contacts with nerve fibers extending from *chata*+ enteric neurons (Fig. 5F and Fig. S5G, H). Analysis of the secretory cell marker 2F11 in *TgBAC(chata:Gal4); Tg(UAS:NTR-mcherry)* animals also revealed a subpopulation of EECs that directly contact *chata*+ enteric nerve fibers (Fig. 5F, Fig. S5I-J and Video 10). Interestingly, those EECs that directly contact *chata*+ enteric neurons include some Trpa1+EECs (Fig. 5G-H). Previous mouse studies demonstrated that some EECs possess neuropods that form synaptic connections with sensory neurons (Bohorquez et al., 2015, Bellono et al., 2017, Kaelberer et al., 2018). We found that zebrafish EECs also possess neuropods marked by the presynaptic marker SV2 (Fig. S5A-C) and are enriched for transcripts encoding presynaptic vesicle proteins (Fig. S5P and Table S1). Similar to mammalian EECs, zebrafish EEC neuropods also contain abundant mitochondria (Fig. S5D) (Bohorquez et al., 2014). Therefore, zebrafish EECs may have an evolutionarily conserved function to signal to neurons, as seen in mammals.

The direct contact of Trpa1+EECs with *chata*+ neurons suggested a direct signal to cholinergic enteric neurons. To investigate whether activation of Trpa1+EECs stimulates *chata*+ enteric neurons, we employed *TgBAC(chata:Gal4); Tg(UAS:Gcamp6s)* zebrafish, which permit recording of *in vivo* calcium activity in *chata*+ neurons (Fig. 5I-J). Upon Trpa1+EEC activation, Gcamp6s fluorescence increased in *chata*+ enteric motor neurons (Fig. 5K, L and Video 11). Immunofluorescence results indicated that Trpa1 is not expressed in *chata*+ enteric motor neurons or in any other ENS cells (Fig. S6O-R). This result indicated that *chata*+ enteric motor neurons cannot be directly activated by optic Trpa1 stimulation but are instead activated via stimulation by Trpa1+ EECs. Previous studies suggested that in addition to the enteric nervous system, efferent vagal nerves also play an important role in modulating intestinal motility (Travagli and Anselmi, 2016). To examine whether Trpa1+EEC induced intestinal motility is indirectly mediated via the vagus nerve, we anatomically disconnected zebrafish intestine from the CNS by decapitation (Fig. S5K). Neither Trpa1+EEC activation-induced intestinal motility nor *chata*+ enteric neuron calcium concentration was affected by decapitation (Fig. S5L-O), suggesting a dispensable role of vagal efferent nerves in Trpa1+EEC induced intestinal motility. These data indicate that Trpa1+EEC induced intestinal motility is mediated by intrinsic enteric circuitry and likely involves *chata*+ enteric neurons.

Previous mouse studies demonstrated that *Trpa1* mRNA is highly enriched in 5-HT-secreting EC cells in the small intestine of mammals (Nozawa et al., 2009). Immunofluorescence staining indicated that, similar to mammals, 5-HT expression in the zebrafish intestinal epithelium is also highly enriched in Trpa1+EECs (Fig. 5M). 5-HT in EECs is synthesized from tryptophan via tryptophan hydroxylase 1 (Tph1) (Li et al., 2011). Zebrafish possess two *Tph1* paralogs, *tph1a* and *tph1b* (Ulhaq and Kishida, 2018), but only *tph1b* is expressed in zebrafish EECs (Fig. S5S). The expression of *tph1b* in Trpa1+EECs was also confirmed by crossing a new *Tg(tph1b:mCherry-NTR)* transgenic line to *TgBAC(trpa1b:EGFP)* zebrafish (Fig. 5N and Fig. S5Q-R, T-U). To investigate whether 5-HT mediates EEC Trpa1-induced intestinal motility, we tested whether a similar response was present in *tph1b*^+/+^ and *tph1b*^-/-^ zebrafish larvae (Tornini et al., 2017) using the Optovin-UV platform. Under baseline conditions, we did not observe a significant difference in intestinal motility between *tph1b*^+/+^ and *tph1b*^-/-^ zebrafish (Fig. S5V). However, in response to UV stimulated EEC Trpa1 activation, intestinal motility was significantly reduced in *tph1b*^-/-^ compared to *tph1b*^+/+^ zebrafish (Fig. 5O). These findings suggest a working model in which Trpa1+EECs signal to cholinergic enteric neurons through 5-HT, which in turn stimulates intestinal motor activity and promotes intestinal motility.

### Chemical and microbial stimulation of EEC Trpa1 signaling activate vagal sensory ganglia

The intestine is innervated by both intrinsic ENS and extrinsic sensory nerves from the brain and spinal cord (Brookes et al., 2013). In mammals, afferent neuronal cell bodies of the vagus nerve reside in the nodose ganglia and travel from the intestine to the brainstem to convey visceral information to the CNS. However, in zebrafish, it is unknown if the vagal sensory system innervates the intestine. The zebrafish vagal sensory ganglia can be labelled using *TgBAC(neurod1:EGFP)* or immunofluorescence staining of the neuronal marker acetylated α Tubulin (Ac-αTub) (Fig. 6B). Using lightsheet confocal imaging, we demonstrated that not only does the vagal ganglia in zebrafish extend projections to the intestine (Fig. 6B-C and Fig. S6A-B) but vagal sensory nerve fibers directly contact a subpopulation of EECs (Fig. 6D). Using the *Tg(neurod1:cre); Tg(β-act2:Brainbow)* transgenic zebrafish system (Gupta and Poss, 2012) (Vagal-Brainbow), in which individual vagal ganglion cells are labeled with different fluorescent colors through Cre recombination (Foglia et al., 2016) (Fig. S6C), we revealed that the zebrafish vagal sensory ganglia cells also directly project to the vagal sensory region in the hindbrain (Fig. 6E-F). Using this Vagal-Brainbow system, we found vagal sensory nerves that are labelled by Cre recombination in both the proximal and distal intestine (Fig. S6D-G). To further visualize the vagal sensory network in zebrafish, we used *Tg(isl1:EGFP)* zebrafish in which EGFP is expressed in vagal sensory ganglia and overlaps with *neurod1* (Fig. 6G and Fig. S6H-J). Our data revealed that after leaving the vagal sensory ganglia, the vagus nerve travels along the esophagus and enters the intestine in the region between the pancreas and the liver (Fig. 6G and Fig. S6I-J). Direct contact of EECs and the vagus nerve could also be observed in *Tg(isl1:EGFP); Tg(neurod1:TagRFP)* zebrafish (Fig. 6H). These data demonstrate the existence of a vagal network in the zebrafish intestine.

**Figure 6.**
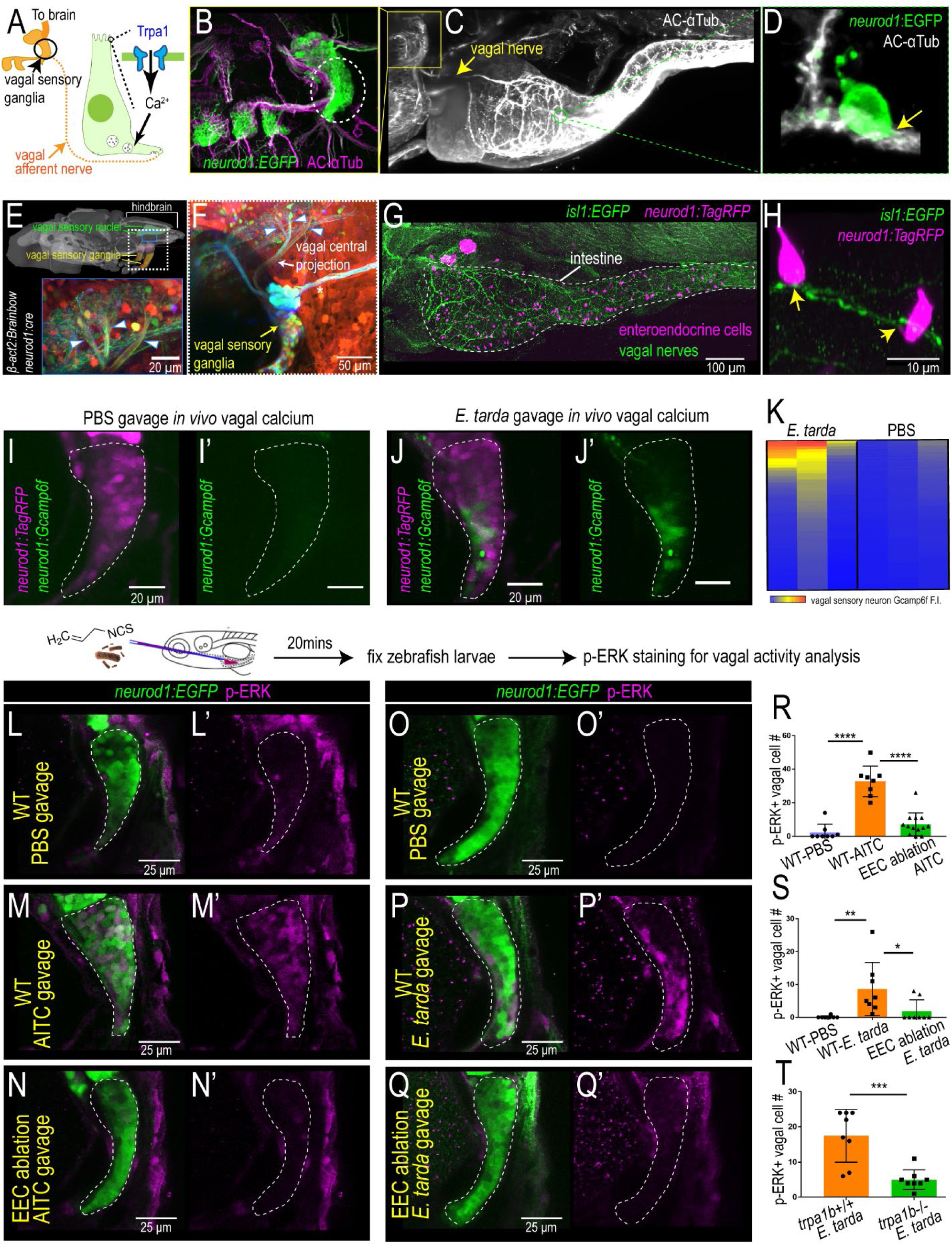
EEC Trpa1 signaling activates vagal sensory ganglia. (A) Working model. (B) Confocal image of zebrafish vagal sensory ganglia labelled with *Tg(neurod1:EGFP)* (green) and acetylated αTubulin antibody staining (AC-αTub, magenta). (C) Lightsheet projection of zebrafish stained acetylated αTubulin antibody. Yellow arrow indicates vagal nerve innervation to the intestine. (D) *neurod1:EGFP*+ EECs (green) directly contact vagal sensory nerve fibers labelled with αTubulin (white). (E) Confocal image of the vagal sensory nucleus in zebrafish larvae hindbrain where - vagal sensory neuron project. Vagal sensory nerve fibers are labeled with different fluorophores through Cre-brainbow recombination in *Tg(neurod1:cre); Tg(βact2:Brainbow)* zebrafish. The 3D zebrafish brain image is generated using mapzebrain (Kunst et al., 2019). (F) Confocal image of vagal sensory ganglia in *Tg(neurod1:cre); Tg(βact2:Brainbow)* zebrafish. Asterisk indicates posterior lateral line afferent nerve fibers. Blue arrowheads indicate three branches from vagal sensory ganglia that project to the hindbrain. (G) Confocal image demonstrates the EEC-vagal network in *Tg(isl1:EGFP); Tg(neurod1:TagRFP)* zebrafish intestine. EECs are labeled as magenta by *neurod1:TagRFP* and the vagus nerve is labeled green by *isl1:EGFP*. (H) EECs (*neurod*1+; magenta) directly contact vagal nerve fibers (*isl1*+; green). Yellow arrows indicate the points where EECs form direct connections with vagal nerve fibers. (I-J) *In vivo* calcium imaging of vagal sensory ganglia in *Tg(neurod1:Gcamp6f); Tg(NBT:DsRed)* zebrafish gavaged with PBS (I) or *E. tarda* (J). (K) Quantification of individual vagal sensory neuron Gcamp6f fluorescence intensity in *E. tarda* or PBS gavaged zebrafish. (L-N) Confocal image of vagal ganglia stained with p-ERK antibody in WT or EEC ablated zebrafish gavaged with PBS or Trpa1 agonist AITC. The vagal sensory ganglia expressing *neurod1:EGFP* are labeled green and activated vagal sensory neurons are labeled magenta by p-ERK antibody staining. (O-Q) Confocal projection of vagal ganglia stained with p-ERK antibody in WT or EEC ablated zebrafish gavaged with PBS or *E. tarda*. (R) Quantification of p-ERK+ vagal sensory neurons in WT or EEC ablated zebrafish following PBS or AITC gavage. (S) Quantification of p-ERK+ vagal sensory neurons in WT or EEC ablated zebrafish following PBS or *E. tarda* gavage. (T) Quantification of p-ERK+ vagal sensory neurons in WT or *trpa1b*-/-zebrafish following *E. tarda* gavage. One-way ANOVA with Tukey’s post test was used in R and S and Student’s t-test was used in T. *p<0.05; **p<0.01; ***p<0.001; ****p<0.0001.

We next investigated whether this vagal network is activated in response to enteric microbial stimulation with *E. tarda*. We gavaged *Tg(neurod1:Gcamp6f); Tg(neurod1:TagRFP)* zebrafish larvae with either PBS or live *E. tarda* bacteria. We found that 30 min after enteric stimulation with Trpa1 agonist AITC or *E. tarda*, but not after PBS vehicle stimulation, Gcamp6f fluorescence intensity significantly increased in a subset of vagal sensory neurons (Fig. 6I-K, Fig. S6K-L and Video 12). This result indicated that acute enteric chemical or microbial stimulation directly activated vagal sensory neurons. To further investigate whether the vagal activation induced by enteric *E. tarda* was mediated by Trpa1+EECs, we used a published method that labels active zebrafish neurons through pERK immunofluorescence staining (Randlett et al., 2015) to measure vagal activity. Delivering AITC into the zebrafish intestine by microgavage (Cocchiaro and Rawls, 2013) increased the number of pERK+ vagal cells compared to PBS treatment (Fig. 6L-N, R). AITC-induced vagal activation was abrogated in the absence of EECs (Fig. 6N, R), indicating that Trpa1 signaling in the intestine increases vagal sensory activity in an EEC-dependent manner. Next, we gavaged live *E. tarda* bacteria into both WT and EEC-ablated zebrafish. Similar to Trpa1 chemical agonist stimulation, *E. tarda* gavage increased the number of activated pERK+ vagal sensory neurons in WT zebrafish (Fig. 6O-Q, S) but not in EEC ablated zebrafish (Fig. 6Q, S). Furthermore, the vagal activation induced by enteric *E. tarda* was dependent on Trpa,1 as pERK+ vagal cell number was significantly reduced in *E. tarda* treated *trpa1b*^-/-^ zebrafish (Fig. 6T). Together, these results reveal that chemical or microbial stimuli in the intestine can stimulate Trpa1+ EECs, which then signal to the vagal sensory ganglia.

### Tryptophan catabolites secreted from *E. tarda* activate the EEC Trpa1 gut-brain pathway

In order to identify the molecular mechanism by which *E. tarda* activates Trpa1 in EECs, we examined the effects of live and killed *E. tarda* cells and cell-free supernatant (CFS) from *E. tarda* cultures on EEC calcium activity (Fig. 7A). Formalin-killed or heat-killed *E. tarda* cells failed to stimulate EECs, however, CFS, at levels comparable to live *E. tarda* cells, stimulated EECs (Fig. 7A-B). The ability of *E. tarda* CFS to activate EECs was diminished in *trpa1b* mutant zebrafish (Fig. 7C), suggesting that a factor secreted from *E. tarda* has the ability to activate Trpa1 in EECs. HPLC-MS analysis revealed that *E. tarda* CFS is enriched for several indole ring-containing tryptophan catabolites (Fig. 7D and Fig. S7A-C), three of the most abundant being indole, tryptophol (IEt), and indole-3-carboxyaldhyde (IAld) (Fig. 7D and Fig. S7A-C). To test if other bacteria secrete tryptophan catabolites like *E. tarda*, we performed similar HPLC-MS analysis of CFS from bacteria we previously tested for EEC activation (Fig.1K). Although several tested bacterial strains produced indole or IAld when cultured in nutrient-rich medium (Fig. S7D), *E. tarda* was the only bacteria that uniquely retained a high level of indole and IAld production when cultured in zebrafish GZM housing water (Fig. 7E), consistent with our finding that *E. tarda* uniquely activates zebrafish EECs when added into GZM water (Fig. 1K). To investigate whether these tryptophan metabolites were directly linked with *E. tarda* pathogenesis, we tested two other *E. tarda* strains (*E. tarda* 23685 and *E. tarda* 15974) which were isolated from human gut microbiota and do not cause fish pathogenesis (Yang et al., 2012, Srinivasa Rao et al., 2003, Nakamura et al., 2013). Both *E. tarda* strains activated EECs and exhibited similar indole and IAld secretion capacity as pathogenic *E. tarda* FL6-60 and *E. tarda* LSE40 (Fig. 7E and Fig. S7G-H). This result suggested that tryptophan catabolites production, EEC Trpa1 activation and its downstream consequences may be distinct from *E. tarda* pathogenesis in fish (Edwardsiellosis), which is characterized by abdominal swelling, petechial hemorrhage in fin and skin, rectal hernia and purulent inflammation in the kidney, liver and spleen (Park et al., 2012).

**Figure 7.**
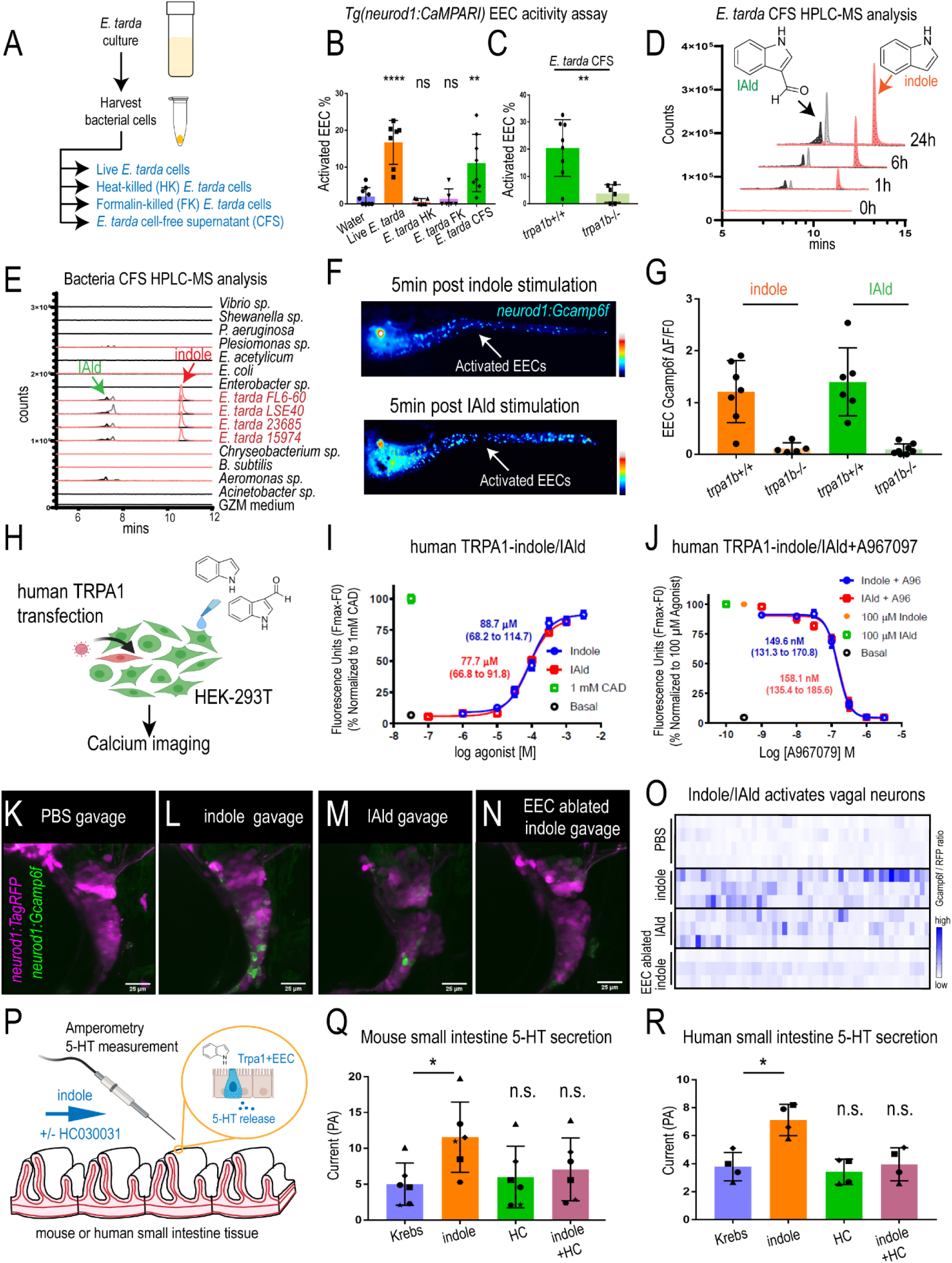
*E. tarda* derived Tryptophan catabolites activate Trpa1 and the EEC-vagal pathway. (A) Method for preparing different fractions from *E. tarda* GZM (zebrafish water) culture. (B) Activated EECs in *Tg(neurod1:CaMPARI)* zebrafish stimulated by different *E. tarda* fractions. (C) Activated EECs in *trpa1b*+/+ and *trpa1b*-/-*Tg(neurod1:CaMPARI)* zebrafish stimulated with *E. tarda* CFS. (D) Screening of supernatants of *E. tarda* in GZM culture medium by HPLC-MS. Samples were collected at 0, 1, 6, 24 h. Abbreviations are as follows: IAld, indole-3-carboxaldehyde; and IEt, tryptophol. Extracted ions were selected for IAld (m/z 145), IEt, (m/z 161), and Indole (m/z 117). (E) Chemical profiles of Trp-Indole derivatives from supernatants of various commensal bacteria in GZM medium for 1 day of cultivation. Values present normalized production of Trp-Indole derivatives based on CFU. (F) *Tg(neurod1:Gcamp6f)* zebrafish stimulated by Indole or IAld. Activated EECs in the intestine are labelled with white arrows. (G) Quantification of EEC Gcamp activity in *trpa1b*+/+ and *trpa1b*-/-*Tg(neurod1:Gcamp6f)* zebrafish stimulated with Indole or IAld. (H) Schematic of experimental design to test effects of indole and IAld on human or mouse Trpa1. (I) Dose-response analysis of the integrated Calcium 6 fluorescence response above baseline (Fmax-F0; maximal change in Ca^2+^ influx) as a function of indole and IAld concentration in human TRPA1 expressing HEK-293T cells. (EC_50_ = 88.7 µM, 68.2-114.7 µM 95% CI for indole; and, EC_50_ = 77.7 µM, 66.8-91.8 µM 95% CI for IAld). Concentration-response data were normalized to 1 mM cinnamaldehyde (CAD), a known TRPA1 agonist. Data represent the mean of 3-4 experiments, each performed with 3-4 replicates. (J) Dose-response analysis of A967079 inhibition of Indole and IAld induced Ca^2+^ influx. (IC_50_ = 149.6 nM, 131.3-170.8 nM 95% CI for Indole; and, IC_50_ = 158.1 nM, 135.4 – 185.6 µM 95% CI for IAld). Concentration-response data of A967079 inhibition was normalized to response elicited by 100 µM agonist (Indole or IAld). (K-N) *In vivo* calcium imaging of vagal sensory ganglia in WT or EEC ablated *Tg(neurod1:Gcamp6f); Tg(neurod1:TagRFP)* zebrafish gavaged with PBS, Indole or IAld. (O) Quantification of individual vagal sensory ganglia cell Gcamp6f fluorescence intensity in WT or EEC ablated zebrafish gavaged with PBS or Indole. (P) Schematic of amperometric measurements to examine the effects of on indole 5-HT secretion in mouse and human small intestinal tissue. (Q) Indole caused a significant increase in 5-HT secretion in mouse duodenum; however, no such effects were observed in the presence of Trpa1 antagonist HC030031. (R) Indole caused a significant increase in 5-HT secretion in human ileum; however, no such effects were observed in the presence of Trpa1 antagonist HC030031. Data in B, C, G, Q, R were presented as mean +/- SD. One-way ANOVA with Tukey’s post test was used in B and Q, Student’s t-test was used in C, H and paired one-way ANOVA with Tukey’s post test was used in P-R. *p<0.05; **p<0.01; ***p<0.001; ****p<0.0001.

We next tested if *E. tarda* tryptophan catabolites activate EECs. Indole and IAld, but not other tested tryptophan catabolites, strongly activated zebrafish EECs in a *trpa1b*-dependent manner (Fig. 7F-G, Fig. S7F and Video 13, 14). Indole and IAld also activated the human TRPA1 receptor transfected into HEK cells (Fig. 7H-J and Fig. S7I-J). Both indole and IAld exhibited full TRPA1 agonist activity with an efficiency comparable to cinnamaldehyde (CAD), a well characterized TRPA1 activator (Fig. 7I-J and Fig. S7I-J) (Macpherson et al., 2007). Both indole and IAld also activated mouse Trpa1, but in a less potent manner (Fig. S7M). Both indole- and IAld-induced human and mouse Trpa1 activation were blocked by the TRPA1 inhibitor A967079 (Fig. 7J and Fig. S7K-L, N). These results establish that indole and IAld that are derived from microbial tryptophan catabolism are novel and evolutionarily-conserved agonists of vertebrate TRPA1 receptors.

Next, we investigated whether indole and IAld can mimic live *E. tarda* bacterial stimulation and activate a similar gut-brain pathway through EEC Trpa1 signaling. Stimulating zebrafish larvae with indole directly induced an increase in intestinal motility (Fig. S8A-D). Using *Tg(neurod1:Gcamp6f);Tg(neurod1:TagRFP)* zebrafish in which vagal calcium levels can be recorded *in vivo*, we found that enteric delivery of indole or IAld by microgavage increased Gcamp6f fluorescence in a subset of vagal sensory neurons (Fig. 7K-O, Fig. S8E-F). This vagal sensory neuron activation induced by enteric indole was abrogated in zebrafish larvae lacking EECs (Fig. 7K-O, Fig. S8E-F). Previous studies demonstrated that the transcription factor Aryl hydrocarbon receptor (AhR) can be activated by many microbially derived tryptophan metabolites including IAld, and gut microbes have been shown to increase intestinal motility via upregulating AhR in mice (Obata et al., 2020, Zelante et al., 2013). To test whether AhR is involved in Trpa1+EEC induced intestinal motility, we applied two well-established AhR inhibitors, CH223191 and folic acid (Kim et al., 2020, Puyskens et al., 2020). However, treatment of zebrafish with these AhR inhibitors was insufficient to block *E. tarda* intestinal accumulation or Optovin-UV-induced intestinal motility (Fig. S8G-L). This finding suggests that *E. tarda* or Trpa1+EEC-induced intestinal motility is not mediated via AhR but instead through a Trpa1+EEC dependent mechanism.

Many microbial tryptophan metabolites, including indole and IAld, are known to be produced by mammalian commensal microbes (Roager and Licht, 2018). Previous studies demonstrated that Trpa1+EECs are restricted to the mouse and human small intestine and not found in the colon (Yang et al., 2019, Billing et al., 2019). We therefore analyzed the tryptophan metabolite concentrations in the small intestine and colon from conventionally-reared mice. We detected several tryptophan metabolites including indole and IAld in the colon as expected, whereas none of these metabolites were detected in the small intestine (Fig. S8G-H). This suggests that these microbial tryptophan metabolites may not significantly contribute to the small intestinal motility under normal physiological condition in mammals. However, under abnormal conditions where microbes and their tryptophan metabolites become elevated in the small intestine (e.g., small intestinal bacterial overgrowth), activated Trpa1+EECs would stimulate enteric and vagal sensory neurons and modulate intestinal motility and brain signaling. To test this model, we used amperometry on fresh tissue sections from human and mouse small intestine to measure the impact of acute indole exposure on 5-HT secretion. As predicted, indole was able to significantly induce 5-HT secretion from both human and mouse small intestine, and this effect was blocked by the Trpa1 inhibitor HC030031 (Fig. 7P-R). These data support our model and further suggest that these microbial tryptophan catabolites may modulate intestinal motility and gut-brain communication in humans.

## DISCUSSION

### Trpa1+EECs are frontline intestinal sensors

To monitor the complex and dynamic chemical and microbial environment within the intestinal lumen, animals evolved specialized sensory cells in the intestinal epithelium known as EECs (Furness et al., 2013). EECs are distinguished from other intestinal epithelial cells by their remarkable ability to respond to a wide range of nutrients and other chemicals and to secrete a variety of peptide hormones and neurotransmitters. Recent studies suggest that mammalian EECs display complex heterogeneity (Gehart et al., 2019). A unique EEC subtype defined in mammals is the enterochromaffin cell (EC) which produces the neurotransmitter 5-HT (Gershon, 2013). A subset of zebrafish EECs are also known to express 5-HT (Roach et al., 2013). In the current study, we identified a Trpa1 expressing EEC subtype that uniquely responds to specific microbial stimulation. We also found that zebrafish Trpa1+EECs include the majority of EECs that express 5-HT, revealing similarity between zebrafish Trpa1^+^EECs and mammalian ECs. In accord, mammalian EC have also been shown to express Trpa1 (Bellono et al., 2017). Our study provides further evidence that Trpa1+EECs respond to chemical and microbial stimuli to inform both the ENS and the vagal sensory nervous system. Thus, Trpa1+EECs appear to be uniquely positioned to protect the organism from harmful chemical and microbial stimuli by regulating GI motility and perhaps sending signals to the brain.

### Microbially derived tryptophan catabolites interact with the host through Trpa1

Trpa1 is a primary nociceptor involved in pain sensation and neuroinflammation. Trpa1 can be activated by several environmental chemical irritants and inflammatory mediators (Bautista et al., 2006), however, it is not known if and how Trpa1 might be activated by microbes. Tryptophan is an essential amino acid that is released in the intestinal lumen by dietary protein digestion or microbial synthesis. It is well known that gut microbes can catabolize tryptophan to produce a variety of metabolites, among which indole was the first discovered and often the most abundant (Smith, 1897). These tryptophan-derived metabolites secreted by gut bacteria can act as interspecies and interkingdom signaling molecules. Some microbially-derived tryptophan catabolites including indole and IAld may regulate host immune homeostasis and intestinal barrier function through ligand binding to the transcription factors, Ahr and Pxr (Venkatesh et al., 2014, Zelante et al., 2013). Another microbial tryptophan catabolite, tryptamine, activates epithelial 5-HT_4_R and increases anion-dependent fluid secretion in the proximal mouse colon (Bhattarai et al., 2018). Though several tryptophan metabolites including IAld can act as AhR agonists (Zelante et al., 2013), conflicting effects of indole on AhR activation have been reported (Heath-Pagliuso et al., 1998, Hubbard et al., 2015, Jin et al., 2014). Whether other host receptors can recognize microbially derived tryptophan catabolites was previously unknown. Here, we present evidence that bacteria-derived tryptophan catabolites activate Trpa1 in zebrafish, human, and mouse. A previous study suggested that indole also activates the yeast TRP channel homolog TRPY1 (John Haynes et al., 2008). This together with our findings point to an ancient role for TRP channels in microbial metabolite sensing. Our results indicate that intestinal colonization by bacteria that produce high levels of tryptophan catabolites (e.g., *E. tarda*) leads to detection of those catabolites by Trpa1+EECs leading to purging of those bacteria by increased intestinal motility. These discoveries were made possible because *E. tarda*, but none of our tested zebrafish commensals, exhibited high capacity to produce and secrete tryptophan catabolites in zebrafish water conditions. Since we did not detect overt pathogenesis in *E. tarda*-treated zebrafish under those experimental conditions, and since many of the *E. tarda* induced responses were recapitulated by indole or IAld alone, we speculate that EEC Trpa1 activation and its downstream consequences reported here are separable from *E. tarda* induced pathogenesis and likely have broader relevance for host-microbial relationships in the gut.

Whereas Trpa1+EECs are abundant in the small intestine of human and rodents (Yang et al., 2019, Nozawa et al., 2009), previous mouse studies demonstrated that Trpa1 in the colon is mainly expressed in mesenchymal cells but not in EECs (Yang et al., 2019, Billing et al., 2019). Conversely to Trpa1+EECs, microbially derived tryptophan metabolites are restricted to the colon and largely absent in small intestine under normal physiological conditions. This is presumably due to the small intestine’s relatively low microbial density or other aspects of digestive physiology that limit microbial production of tryptophan metabolites (Kastl et al., 2020). This suggests that microbially derived tryptophan metabolites do not activate Trpa1+EECs to affect intestinal motility under normal physiological conditions. On one hand, Trpa1+EECs may act as a reserve host protective mechanism that detects tryptophan catabolites accumulating due to aberrant overgrowth of small intestinal microbiota or invasion of specific commensals or pathogens like *E. tarda* that precociously produce those catabolites, and in response increases intestinal motility to purge that particular community. Loss or impairment of that protective mechanism may result in overgrowth or dysbiosis of small intestinal microbial communities or an increased risk of enteric infection. On the other hand, excessive or chronic activation of those Trpa1+EECs may result in pathophysiological changes. One such scenario may be small intestinal bacteria overgrowth (SIBO), which is prevalent in patients suffering from diarrhea-dominant irritable bowel syndrome (IBS) (Ghoshal et al., 2017). IBS is a complicated disease that display comorbidities of both impairment of GI motility and CNS symptoms. The cause of SIBO in IBS is incompletely understood although several studies demonstrated that some of the indole producing bacteria like *Escherichia* coli exhibit high abundance in the small intestine of SIBO associated IBS patients (Ghoshal et al., 2014, Leite et al., 2020, Avelar Rodriguez et al., 2019). Our findings raise the possibility that SIBO leads to an increase of microbial tryptophan metabolite production in the small intestine, which then activates Trpa1+EECs to increases intestinal motility and modulate CNS activity through the vagal nerve, resulting in the complex comorbidities of intestinal and psychiatric disorders in IBS.

### Gut microbiota-EEC-ENS communication

Nerve fibers do not penetrate the gut epithelium therefore, sensation is believed to be a transepithelial phenomenon as the host senses gut contents through the relay of information from EECs to the ENS (Gershon, 2004). Using an *in vitro* preparation of mucosa–submucosa, mechanical or electrical stimulation of mucosa was shown to activate submucosal neuronal ganglia, an effect blocked by a 5-HT_1_R antagonist (Pan and Gershon, 2000). Consistent with these previous findings, our zebrafish data suggest a model that 5-HT released from Trpa1+EECs stimulates intrinsic primary afferent neurons (IPANs) which then activate secondary neurons to promote intestinal motility through the local enteric EEC-ENS circuitry.

90% of 5-HT in the intestine is produced by EC cells, and therefore, EC cell 5-HT secretion was thought to be important in regulating intestinal motility (Gershon, 2013). This hypothesis, however, was challenged by recent findings that depletion of EC 5-HT production in *Tph1*^-/-^ mice had only minor effects on gastric emptying, intestinal transit, and colonic motility (Li et al., 2011). Therefore, the physiological role of EC 5-HT production and secretion remains unclear. Our data suggest that EEC 5-HT production may be necessary for intestinal motility changes in response to environmental chemical or microbial stimuli, but not for intestinal motility under normal physiological conditions. Mice raised germ free displayed lower 5-HT content in the colon, however no significant difference of 5-HT production was observed in the small intestine compared to colonized mice (Yano et al., 2015). Whether gut microbiota regulate small intestinal 5-HT secretion and signaling remains unknown. Our data suggest a model in which specific microbial communities or individual microbial types may stimulate 5-HT secretion from Trpa1+EECs to modulate small intestinal motility by producing tryptophan catabolites. This may provide a new mechanism by which gut microbiota can regulate 5-HT signaling in the small intestine.

Gut microbiota are believed to be important for regulating GI motility. Mice raised under germ-free conditions display longer intestinal transit time and abnormal colonic motility (Vincent et al., 2018). In humans, the association between gut microbiota and GI motility is also evident, especially in IBS. IBS patients usually display alterations in GI motility without intestinal inflammation and many patients with diarrhea-predominant IBS developed their symptoms after an acute enteric bacterial infection (e.g., post-infectious IBS) and displayed alterations in gut microbiota composition (Ghoshal and Gwee, 2017). The mechanisms underlying gut microbiota-induced GI motility and the development of pathological conditions like IBS are unclear. Here, we provide evidence that a specific set of microbially derived tryptophan catabolites directly stimulate intestinal motility by activating EEC-ENS signaling via release of 5-HT. Indole, IAld and other tryptophan catabolites are produced by a wide range of gut bacteria, so we expect our results to be applicable to commensal and pathogenic bacteria and their host interactions. These findings offer new possibilities for treatment of gut microbiota associated GI disorders, by targeting microbial tryptophan catabolism pathways, microbial or host degradation of those catabolites, or targeting EEC microbial sensing and EEC-ENS signaling pathways.

### Gut microbiota-EEC-CNS communication

The vagus nerve is the primary sensory pathway by which visceral information is transmitted to the CNS. Recent evidence suggests that the vagus nerve may play a role in communicating gut microbial information to the brain (Fulling et al., 2019, Breit et al., 2018, Bonaz et al., 2018). For example, the beneficial effects of *Bifidobacterium longum* and *Lactobacillus rhamnosus* in neurogenesis and behavior were abolished following vagotomy (Bercik et al., 2011, Bravo et al., 2011). However, direct evidence for whether and how vagal sensory neurons perceive and respond to gut bacteria has been lacking. Our results demonstrate that both live *E. tarda* and *E. tarda*-derived tryptophan catabolites activate vagal sensory ganglia through EEC Trpa1 signaling. Previous findings have shown that EC cells transmit microbial metabolite and chemical irritant stimuli to pelvic fibers from the spinal cord dorsal root ganglion (Bellono et al., 2017). Our findings here demonstrate that, in addition to spinal sensory nerves, EEC-vagal signaling is an important pathway for transmitting specific gut microbial signals to the CNS. The vagal ganglia project directly onto the hindbrain, and that vagal-hindbrain pathway has key roles in appetite and metabolic regulation (Grill and Hayes, 2009, Han et al., 2018, Travagli et al., 2006, Berthoud et al., 2006). Our findings raise the possibility that certain tryptophan catabolites, including indole, may directly impact these processes as well as emotional behavior and cognitive function (Jaglin et al., 2018). If so, this pathway could be manipulated to treat gut microbiota-associated neurological disorders.

## Supporting information

Key Resources Table

Table S1_zebrafish EEC RNAseq

Table S2_Compare zebrafish and mammalian EECs

Table S3_Specific EEC genes

## ACKNOWLEDGEMENTS

This work was supported by NIH R01-DK093399, R01-DK109368, VA-BX002230, and a Pew Scholars Innovation Award from the Pew Charitable Trusts. S.V.J. and S.-E.J. were supported by cooperative agreement U01ES030672 of the NIH CounterACT Program. K.D.P. was supported by R01 GM074057 and R35 HL150713. L.Y. was supported by NIH T32-DK007568. C. D.C. was supported by NIH F32AT010415. D.J.K. was supported by the Australian Research Council, DP190103525. The content is solely the responsibility of the authors and does not necessarily represent the views of the NIH.

## AUTHOR CONTRIBUTIONS

Conceptualization: L.Y., R.L., J.R.; Formal Analysis: L.Y., S.J., M.B., C.C., C.L.; Funding acquisition: K.P., D.J., S.-E.J., R.L., J.R.; Investigation: L.Y., S.J., M.B., C.C., D.T., A.M., H.L., J. C., D.K.; Methodology: L.Y., S.J., M.B., C.C., D.T., A.M., D.K.; Resources: L.Y., J. W., J.T., K.P.; Supervision: J.C., R.L., J.R.; Visualization: L.Y., S.J., M.B.; Writing – original draft: L.Y.; Writing – review & editing: L.Y., R.L., J.R., K.P., C.L..

## STAR METHODS

### Resource Availability

#### Lead Contact

Further information and requests for resources and reagents should be directed to and will be fulfilled by the Lead Contact, John F. Rawls.

### Materials Availability

Zebrafish strains and plasmids generated in this study are available upon request from the Lead Contact.

### Data and Code Availability

Sequencing reads generated as part of this study are available at Gene Expression Omnibus accession GSE151711.

### Experimental Model and Subject Details

#### Zebrafish strains and husbandry

All zebrafish experiments conformed to the US Public Health Service Policy on Humane Care and Use of Laboratory Animals, using protocol numbers A115-16-05 and A096-19-04 approved by the Institutional Animal Care and Use Committee of Duke University. For experiments involving conventionally raised zebrafish larvae, adults were bred naturally in system water and fertilized eggs were transferred to 100mm petri dishes containing ∼25 mL of egg water at approximately 6 hours post-fertilization. The resulting larvae were raised under a 14 h light/10 h dark cycle in an air incubator at 28°C at a density of 2 larvae/mL water. All the experiments performed in this study ended at 6 dpf unless specifically indicated. The strains used in this study are listed in Key Resources Table. All lines were maintained on a mixed Ekkwill (EKW) background.

### Bacterial strains and growing conditions

All bacterial strains in this study were cultured at 30°C in Trypticase soy broth (TSB) or Gnotobiotic zebrafish medium (GZM) (Pham et al., 2008). Tryptic Soy Agar (TSA) plate was used for streaking bacterial from glycerol stock or performing colony forming unit (CFU) experiments. The antibiotic carbenicillin was used to select *E. tarda* LSE that express mCherry at the working concentration of 100 µg/mL.

### Method Details

#### Generating transgenic zebrafish

The Gateway Tol2 cloning approach was used to generate the *neurod1:CaMPARI* and *neurod1:cre* plasmids (Kawakami, 2007, Kwan et al., 2007). The 5kb pDONR-neurod1 P5E promoter was previously reported (McGraw et al., 2012) and generously provided by Dr. Hillary McGraw. The pME-cre plasmid as reported previously (Cronan et al., 2016) was generously donated by Dr. Mark Cronan. The pcDNA3-CaMPARI plasmid was reported previously (Fosque et al., 2015) and obtained from Addgene. The CaMPARI gene was cloned into pDONR-221 plasmid using BP clonase (Invitrogen, 11789-020) to generate PME-CaMPARI. pDONR-neurod1 P5E and PME-CaMPARI were cloned into pDestTol2pA2 using LR Clonase (ThermoFisher,11791). Similarly, pDONR-neurod1 P5E and pME-cre were cloned into pDestTol2CG2 containing a cmlc2:EGFP marker. The final plasmid was sequenced and injected into the wild-type EKW zebrafish strain and the F2 generation of alleles *Tg(neurod1:CaMPARI)*^rdu78^ and *Tg(neurod1:cre; cmlcl2:EGFP)*^rdu79^ were used for this study.

To make transgenic lines, that permit specific EEC ablation, we used *Tg(neurod1:cre)* and *TgBAC(gata5:loxp-mCherry-stop-loxp-DTA)* new transgenic system. This system consists of two new transgene alleles - one expressing Cre recombinase from the *neurod1* promoter (in EECs, CNS, and islets) and a second expressing the diphtheria toxin (DTA) in *gata5*+ cells (in EECs, other IECs, heart, and perhaps other cell types) only in the presence of Cre (Fig. 3F). As the only cells known to co-express *neurod1* and *gata5* in the zebrafish larvae, EECs are ablated whereas non-EEC cell populations, including islets and the CNS, remain unaffected (Fig. 3G). A small percentage of EECs remained in the distal intestine presumably due to the low level of *gata5* expression in that region (Fig. S4C). The method for generating *Tg(neurod1:cre)* was described above. To generate the *TgBAC(gata5:loxp-mCherry-stop-loxp-DTA)* transgenic line, the translational start codon of *gata5* in the BAC clone DKEYP-73A2 was replaced with the loxP-mCherry-STOP-loxP-DTA (RSD) cassette by Red/ET recombineering technology (GeneBridges). For recombination with arms flanking the RSD cassette, the 5’ homologous arm used was a 716 bp fragment upstream of the start codon and the 3’ homologous arm was a 517 bp downstream fragment. The vector-derived loxP site was replaced with an I-SceI site using the same technology. The final BAC was purified using the Qiagen Midipre kit, and coinjected with I-SceI into one-cell stage zebrafish embryos. The full name of this transgenic line is *Tg(gata5:loxP-mCherry-STOP-loxP-DTA)*^*pd315*^.

*Tg(tph1b:mCherry-NTR)*^*pd275*^ zebrafish were generated using I-SceI transgenesis in an Ekkwill (EK) background. Golden Gate Cloning with BsaI-HF restriction enzyme (NEB) and T4 DNA ligase (NEB) was used to generate the tph1b:mCherry-NTR plasmid by cloning the 5kb *tph1b* promoter sequence (tph1bP GG F: GGTCTCGATCGGtctaaggtgaatctgtcacattc; tph1bP GG R: GGTCTCGGCTACggatggatgctcttgttttatag), mCherry (mC GG F: GGTCTCGTAGCC gccgccaccatggtgag; mC GG2 R: GGTCTCGGTACCcttgtacagctcgtccatgccgcc), a P2A polycistronic sequence and triple mutant variant nitroreductase (Mathias et al., 2014) (mutNTR GG F: GGTCTCGGTACCtacttgtacaagggaagcggagc; mutNTR GG2 R: GGTCTCCCATGC caggatcggtcgtgctcga), into a pENT7 vector backbone with a poly-A tail and I-SceI sites (pENT7 mCN GG F: GGTCTCGCATGGacacctccccctgaacctg; pENT7 mCN GG R: GGTCTCCCGATC gtcaaaggtttggggtccgc). 500 pL of 25 ng/μL plasmid, 333 U/mL I-SceI (NEB), 1x I-SceI buffer, 0.05% Phenol Red (Sigma-Aldrich) solution was injected into EK 1-cell zebrafish embryos. F0 founders were discovered by screening for fluorescence in outcrossed F1 embryos.

### RNA sequencing and bioinformatic analysis

To isolate zebrafish EECs and other IECs, we crossed two transgenic zebrafish lines, one that specifically expresses enhanced green fluorescent protein (EGFP) in all intestinal epithelial cells (*TgBAC(cldn15la:EGFP)*) (Alvers et al., 2014) and a second that expresses red fluorescent protein (RFP) in EECs, pancreatic islets, and the central nervous system (CNS) (*Tg(neurod1:TagRFP)*) (McGraw et al., 2012). The FACS-isolated EECs were identified by *cldn15la:EGFP*+; *neurod1: TagRFP*+; and the other IECs were identified by *cldn15la:EGFP*+; *neurod1:TagRFP*-. Conventionalized (CV) and germ-free (GF) *TgBAC(cldn15la:EGFP); Tg(neurod1:TagRFP)* ZM000 fed zebrafish larvae were derived and reared using our published protocol (Pham et al., 2008) for Flow Activated Cell Sorting (FACS) to isolate zebrafish EECs and other IECs. The protocol for FACS was adopted from a previous publication (Espenschied et al., 2019). Replicate pools of 50-100 double transgenic *TgBAC(cldn15la:EGFP); Tg(neurod1:TagRFP)* zebrafish larvae were euthanized with Tricaine and washed with deyolking buffer (55 mM NaCl, 1.8 mM KCl and 1.25 mM NaHCO_3_) before they were transferred to dissociation buffer [HBSS supplemented with 5% heat-inactivated fetal bovine serum (HI-FBS, Sigma, F2442) and 10 mM HEPES (Gibco, 15630–080)]. Larvae were dissociated using a combination of enzymatic disruption using Liberase (Roche, 05 401 119 001, 5 μg/mL final), DNaseI (Sigma, D4513, 2 μg/mL final), Hyaluronidase (Sigma, H3506, 6 U/mL final) and Collagenase XI (Sigma, C7657, 12.5 U/mL final) and mechanical disruption using a gentleMACS dissociator (Miltenyi Biotec, 130-093-235). 400 μL of ice-cold 120 mM EDTA (in 1x PBS) wwas added to each sample at the end of the dissociation process to stop the enzymatic digestion. Following addition of 10 mL Buffer 2 [HBSS supplemented with 5% HI-FBS, 10 mM HEPES and 2 mM EDTA], samples were filtered through 30 μm cell strainers (Miltenyi Biotec, 130-098-458). Samples were then centrifuged at 1800 rcf for 15 minutes at room temperature. The supernatant was decanted, and cell pellets were resuspended in 500 μL Buffer 2. FACS was performed with a MoFlo XDP cell sorter (Beckman Coulter) at the Duke Cancer Institute Flow Cytometry Shared Resource. Single-color control samples were used for compensation and gating. Viable EECs or IECs were identified as 7-AAD negative.

Samples from three independent experimental replicates were performed. 250-580 EECs (n=3 for each CV and GF group) and 100 IECs (n=3 for each CV and GF group) from each experiment were used for library generation and RNA sequencing. Total RNA was extracted from cell pellets using the Argencourt RNAdvance Cell V2 kit (Beckman) following the manufacturer’s instructions. RNA amplification prior to library preparation had to be performed. The Clontech SMART-Seq v4 Ultra Low Input RNA Kit (Takara) was used to generate full-length cDNA. mRNA transcripts were converted into cDNA through Clontech’s oligo(dT)-priming method. Full length cDNA was then converted into an Illumina sequencing library using the Kapa Hyper Prep kit (Roche). In brief, cDNA was sheared using a Covaris instrument to produce fragments of about 300 bp in length. Illumina sequencing adapters were then ligated to both ends of the 300bp fragments prior to final library amplification. Each library was uniquely indexed allowing for multiple samples to be pooled and sequenced on two lanes of an Illumina HiSeq 4000 flow cell. Each HiSeq 4000 lane could generate >330M 50bp single end reads per lane. This pooling strategy generated enough sequencing depth (∼55M reads per sample) for estimating differential expression. Sample preparation and sequencing was performed at the GCB Sequencing and Genomic Technologies Shared Resource.

Zebrafish RNA-seq reads were mapped to the danRer10 genome using HISAT2(Galaxy Version 2.0.5.1) using default settings. Normalized counts and pairwise differentiation analysis were carried out via DESeq2 (Love et al., 2014) with the web based-galaxy platform: https://usegalaxy.org/. For the purpose of this study, we only displayed the CV EEC (n=3) and CV IEC (n=3) comparison and analysis in the Results section. The default significance threshold of FDR < 5% was used for comparison. Hierarchical clustering of replicates and a gene expression heat map of RNA-seq data were generated using the online expression heatmap tool: http://heatmapper.ca/expression/. The human and mouse RNA-seq raw counts data were obtained from the NCBI GEO repository: human, GSE114853; mouse, GSE114913 (Roberts et al., 2019). Pairwise differentiation analysis of human jejunum CHGA+ (n=11) and CHGA-(n=11) and mouse duodenum Neurod1+ (n=3) and Neurod1-(n=3) was performed using DESeq2. The mouse and zebrafish ortholog Gene ID conversion was downloaded from Ensemble. The genes that were significantly enriched (P_FDR_<0.05) in the human and mouse EEC data sets were used to query the zebrafish EEC RNA seq dataset and data were plotted using Graphpad Prism7. RNA-seq data generated in this study can be accessed under Gene Expression Omnibus accession GSE151711.

### Recording *in vivo* EEC activity

CaMPARI undergoes permanent green-to-red photoconversion (PC) under 405 nm light when calcium is present. This permanent conversion records the calcium activity for all areas illuminated by PC-light. Red fluorescence intensity correlates with calcium activity during photoconversion (Fosque et al., 2015). In the *Tg(neurod1:CaMPARI)* zebrafish line, the CaMPARI (calcium-modulated photoactivatable ratiometric integrator) transgene is expressed under control of the -5kb promoter cloned from the zebrafish *neurod1* locus. CaMPARI mRNA is transcribed and the CaMPARI protein is expressed in cells that are able to activate the *neurod1* promoter. There are multiple cell types in the zebrafish body that are sufficient to activate the *neurod1* promoter, including all EECs in the intestine (Ye et al., 2019). CaMPARI protein is a calcium indicator protein that binds calcium and converts from green fluorescence to red fluorescence in the presence of UV light. This protein is engineered and described in detail in a previous publication (Fosque et al., 2015). We use this transgenic model to measure the level of intracellular calcium in EECs. Similar to neurons, it is well known that when extracellular stimulants act on various receptors on EECs, this leads to an increase of intracellular calcium either due to calcium influx through calcium channels in the plasma membrane or release of calcium stored in the ER. Through either of these pathways, increased intracellular calcium then directly triggers EECs to release hormone/neurotransmitter vesicles. To record *in vivo* EEC activity using the CaMPARI platform, conventionally raised *Tg(neurod1:CaMPARI)* zebrafish larvae were sorted at 3 dpf and maintained in Gnotobiotic Zebrafish Media (GZM) (Pham et al., 2008) with 1 larvae/mL density. At 6 dpf, for each experimental group, ∼20 larvae were transferred into 50mL conical tubes in 2 mL GZM medium. The larvae were adjusted to the new environment for 30 mins before stimuli were added to each conical tube. For nutrient stimulation, since linoleate, oleate and laurate are not soluble in water, a bovine serum albumin (BSA) conjugated fatty acid solution was generated as described previously (Ye et al., 2019). 2 mL linoleate, oleate, laurate, butyrate or glucose was added to the testing tube containing ∼20 zebrafish larvae in 3 mL GZM. The final stimulant concentrations were: linoleate (1.66 mM), oleate oleate (1.66 mM), laurate (1.66 mM), butyrate (2 mM) and glucose (500 mM). Zebrafish larvae were stimulated for 2 mins (fatty acids) or 5 mins (glucose) before the UV pulse. For bacterial stimulation, single colonies of the different bacterial strains were cultured aerobically in tryptic soy broth (TSB) at 30°C overnight (rotating 50-60 rpm, Thermo Fisher Tissue Culture Rotator CEL-GRO #1640Q)(see strains listed in Key Resources Table). O/N TSB cultured bacteria were harvested, washed with GZM and resuspended in 2 mL GZM. 2 mL bacteria were then added to a test tube containing ∼20 zebrafish larvae in 3 mL GZM. The final concentration of the bacterial is ∼ 10^8^ CFU/ml. Zebrafish were then stimulated for 20mins before treated with a UV pulse. A customized LED light source (400 nm-405 nm, Hongke Lighting CO. LTD) was used to deliver a UV light pulse (100 W power, DC32-34 V and 3500 mA) for 30 seconds. Following the UV pulse, zebrafish larvae were transferred to 6-well plates. To block spontaneous intestinal motility and facilitate *in vivo* imaging, zebrafish larvae were incubated in 20 µM 4-DAMP (mAChR blocker), 10 µM atropine (mAChR blocker) and 20 µM clozapine (5-HTR blocker) for 30 mins. Zebrafish larvae were then anesthetized with Tricaine (1.64 mg/ml) and mounted in 1% low melting agarose and imaged using a 780 Zeiss upright confocal microscope in the Duke Light Microscope Core Facility. Z-stack confocal images were taken of the mid-intestinal region in individual zebrafish. The laser intensity and gain were set to be consistent across different experimental groups. The resulting images were then processed and analyzed using FIJI software (Schindelin et al., 2012). To quantify the number of activated EECs, the color threshold was set for the CaMPARI red channel. EECs surpassing the color threshold were counted as activated EECs. The CaMPARI green channel was used to quantify the total number of EECs in each sample. The ratio of activated EECs to the total EEC number was calculated as the percentage of activated EECs. As reported in Fig. S1A-F, in *Tg(neurod1:CaMPARI)* zebrafish model, in addition to EECs, CaMPARI is also expressed in other *neurod1*+ cells including CNS and pancreatic islet. Therefore, the *Tg(neurod1:CaMPARI)* model can also be used to measure the activity of the CNS and pancreatic islet. However, the method we described above permit us to specifically analyze EEC signal through restricting our image inquiry in the middle intestine, a region in which only EECs express CaMPARI.

To record *in vivo* EEC activity using the *Tg(neurod1:Gcamp6f)* system, we used our published protocol with slight modification (Ye et al., 2019). In brief, unanesthetized zebrafish larvae were gently mounted in 3% methylcellulose. Excess water was removed and zebrafish larvae were gently positioned with right side up. Zebrafish were then moved onto an upright Leica M205 FA fluorescence stereomicroscope equipped with a Leica DFC 365FX camera. The zebrafish larvae were allowed to recover for 2mins before 100 µL of test agent was pipetted directly in front of the mouth region. Images were then recorded every 10 seconds. The stimulants used in this study are listed in Supplemental Table 1. The data shown in Fig. 2O-R, depicting the EEC responses to *E. tarda* stimulation, were obtained by mounting unanesthetized zebrafish larvae in 1% low melting agarose. A window (5 × 5 mm) was cut to expose the head region of the zebrafish. 10 µL of *E. tarda* culture [∼10^9^ Colony Forming Unit (CFU)] were delivered at the zebrafish mouth area. Images were recorded every 10 secs for 20 mins. Image processing and analysis were performed using FIJI software. Time-lapse fluorescence images were first aligned to correct for experimental drift using the plugin “align slices in stack.” Normalized correlation coefficient matching and bilinear interpolation methods for subpixel translation were used for aligning slices (Tseng et al., 2012). The plugin “rolling ball background subtraction” with the rolling ball radius=10 pixels was used to remove the large spatial variation of background intensities. The Gcamp6f fluorescence intensity in the intestinal region was then calculated for each time point. The ratio of maximum fluorescence (Fmax) and the initial fluorescence (F0) was used to measure EEC calcium responses.

### Immunofluorescence staining and imaging

Whole mount immunofluorescence staining was performed as previously described (Ye et al., 2019). In brief, ice cold 2.5% formalin was used to fix zebrafish larvae overnight at 4°C. The samples were then washed with PT solution (PBS+0.75%Triton-100). The skin and remaining yolk were then removed using forceps under a dissecting microscope. The deyolked samples were then permeabilized with methanol for more than 2 hrs at -20°C. Samples were then blocked with 4% BSA at room temperature for more than 1 hr. The primary antibody was diluted in PT solution and incubated at 4°C for more than 24 hrs. Following primary antibody incubation, the samples were washed with PT solution and incubated overnight with secondary antibody with Hoechst 33342 for DNA staining. Imaging was performed with Zeiss 780 inverted confocal and Zeiss 710 inverted confocal microscopes with 40× oil lens. The primary antibodies were listed in Supplemental Table 1. The secondary antibodies in this study were from Alexa Fluor Invitrogen were used at a dilution of 1:250.

To quantify vagal activity by pERK staining, we used a published protocol with slight modification (Randlett et al., 2015). Zebrafish larvae were quickly collected by funneling through a 0.75 mm cell strainer and dropped into a 5mL petri dish containing ice cold fix buffer (2.5% formalin+ 0.25% Triton 100). Larvae were fixed overnight at 4°C, then washed 3 times in PT (PBS+ 0.3% Triton 100), treated with 150 mM Tris-HCl (PH=9) for 15 mins at 70°C, washed with PT and digested with 0.05% trypsin-EDTA on ice for 45 mins. Following digestion, samples were then washed with PT and transferred into block solution [PT + 1% bovine serum albumin (BSA, Fisher) + 2% normal goat serum (NGS, Sigma) + 1% dimethyl sulfoxide (DMSO)]. The primary antibodies [pERK (Cell signaling); tERK (Cell signaling); GFP (Aves Lab)] were diluted in block solution (1:150 for pERK; 1:150 for tERK and 1:500 for GFP) and samples were incubated in 100 µl of primary antibody overnight at 4°C. Following primary antibody incubation, samples were then washed with PT and incubated with secondary antibody overnight at 4°C. Samples were then washed with PBS, mounted in 1% LMA and imaged using a Zeiss 780 upright confocal microscope.

### Zebrafish *E. tarda* colonization

For *E. tarda* colonization experiments, fertilized zebrafish eggs were collected, sorted and transferred into a cell culture flask containing 80 mL GZM at 0 dpf. At 3 dpf, dead embryos and 60 mL GZM were removed and replaced with 50 mL fresh GZM in each flask. To facilitate consistent commensal gut bacterial colonization, an additional 10 mL of filtered system water (5 μm filter, SLSV025LS, Millipore) were added to each flask. Overnight *E. tarda* mCherry (Amp^r^, see details in Supplemental Table 1) culture was harvested, washed three times with GZM. 150 µL of GZM-washed *E. tarda* mCherry culture were inoculated into each flask. The *E. tarda* concentration is ∼10^6^ CFU/ml. Daily water changes (60 ml) was performed and 200 µL autoclaved solution of ZM000 food (ZM Ltd.) was added from 3 dpf to 6 dpf as previously described (Pham et al., 2008). At 6 dpf, zebrafish larvae were subjected to fluorescence imaging analysis or CFU quantification. For fluorescence imaging analysis, zebrafish larvae were anesthetized with Tricaine (1.64 mg/ml), mounted in 3% methylcellulose and imaged with a Leica M205 FA upright fluorescence stereomicroscope equipped with a Leica DFC 365FX camera. For CFU quantification, digestive tracts were dissected and transferred into 1 mL sterile PBS which was then mechanically disassociated using a Tissue-Tearor (BioSpec Products, 985370). 100 µL of serially diluted solution was then spread on a Tryptic soy agar (TSA) plate with Carbenicillin (100 µg/ml) and cultured overnight at 30°C under aerobic conditions. The mCherry+ colonies were quantified from each plate and *E. tarda* colony forming units (CFUs) per fish were calculated.

### Zebrafish microgavage and chemical treatment

For delivering bacterial or chemicals specifically to the intestine, we adopted our established microgavage technique (Cocchiaro and Rawls, 2013). Zebrafish were anesthetized with 1 mg/mL α-Bungarotoxin (α-BTX) and the gavage procedure was performed as previously described using microinjection station (Cocchiaro and Rawls, 2013). For bacteria gavage experiments, 1ml of overnight bacterial culture was harvested, pelleted, washed with PBS and resuspended in 100ul PBS. ∼ 8nl was then delivered into the zebrafish intestine using microgavage. The gavaged zebrafish was then transferred into egg water and mounted in 1% LMA for imaging. For chemical gavage experiments, ∼ 8nl of AITC (100mM), indole (1mM) and IAld (1mM) was gavaged into the intestine.

For Trpa1 inhibition, Trpa1 antagonist HC030031 (280µM) was treated 2 hours before and during the 30 mins of *E. tarda* stimulation. For AhR inhibition, two AhR inhibitors, CH030031 and folic acid, were selected based on previous publications (Puyskens et al., 2020, Kim et al., 2020). CH030031 is a well-established specific AhR inhibitor (Choi et al., 2012). Whereas folic acid is shown to act as a competitive AhR antagonist at the concentration as low as 10ng/ml (Kim et al., 2020). For the *E. tarda* treatment experiment, DMSO, CH030031 (1µM) or folic acid (10µM) was added into zebrafish water at 3 dpf zebrafish at the same time as *E. tarda* administration. The AhR inhibitors were replenished during daily water changes, and zebrafish were analyzed at 6 dpf. For the Optovin-UV experiment, overnight Optovin treated zebrafish were treated for 2 hours with DMSO, CH030031 (10µM) or folic acid (10µM). As demonstrated by previous study, 2-hour 10µM and 3-day 0.5µM CH030031 treatment is sufficient to inhibit larve zebrafish AhR signaling (Puyskens et al., 2020, Sun et al., 2019b, Yue et al., 2017). The concentration of FA was chosen based on zebrafish tolerance and a previous study shown the treatment of early zebrafish embryos with 0.05µM FA inhibits AhR signaling (Yue et al., 2017).

### Optic EEC activation

For EEC Trpa1 activation using the Optovin platform, zebrafish larvae were treated with 10 µM Optovin overnight. Following Optovin treatment, unanesthetized zebrafish were mounted in 1% LMA and imaged under a 780 upright Zeiss confocal microscope using 20× water objective lenses. For all the experiments, the mid-intestine region was imaged (Fig. S5D). The intestinal epithelium was selected as the region of interest (ROI) (Fig. S5A). Serial images were obtained at 1 s/frame. A 405 nm pulse of light was applied to the ROI at 1 pulse/10s. For some experiments (Fig. 4D-F, Fig. S5B-G), the images were obtained at 10s/frame. When measuring Optovin effects on intestinal motility in *ret*-/-, *sox10*-/- or *tph1b*-/-zebrafish larvae, embryos were collected from heterozygous zebrafish. *ret*-/-zebrafish were identified by lack of ENS and deflated swim bladder (Knight et al., 2011), *sox10*-/-zebrafish were identified by lack of pigment (Rolig et al., 2017), and *tph1b*-/-zebrafish were identified by PCR-based genotyping (Tornini et al., 2017).

Photoactivation of channelrhodopsin (ChR2) in EECs was performed in *Tg(neurod1:Gal4, cmlc2;EGFP); Tg(UAS:ChR2-mCherry)* transgenic zebrafish. In this model, ChR2 expression in EECs is mosaic. At 6 dpf, unanesthetized zebrafish larvae were mounted in 1% LMA. Photoactivation and imaging were performed with a Zeiss 780 upright confocal using 20× water objective lenses. Individual ChR2+ EECs were selected as ROI (Fig. S5H, I). Serial images were obtained at 1 s/frame. The 488 nm and 458 nm pulses were applied to the selected ROI at 1 pulse/s. For selectively activating *trpa1b*+ or *trpa1b*-ChR2 expressing EECs, *Tg(neurod1:Gal4, cmlc2;EGFP); Tg(UAS:ChR2-mCherry)* was crossed with *TgBAC(trpa1b:EGFP)*. In each zebrafish, either Trpa1+ChR2+ EEC or Trpa1-ChR+ EEC was selected to activate and examine the motility pre and post activation. Each dot in updated Fig. 4L and Fig. S3H represent data from individual zebrafish. A snapshot of the intestinal area was obtained to determine the *trpa1b*+ChR2+ and *trpa1b*-ChR2+ EECs (Fig. S5H, I) and light pulses were applied to the selected EECs as indicated above. Due to the mosaic expression of ChR2 in the EECs in the Gal4-UAS transgenic system, the ChR2^+^EECs in both proximal and middle intestinal regions are selected.

To determine whether Optovin-UV or ChR2 was sufficient to activate EECs, *Tg(neurod1:Gcamp6f)* zebrafish were used. To facilitate EEC calcium imaging under the confocal microscope, zebrafish larvae were incubated in 20 µM 4-DAMP, 10 µM atropine and 20 µM clozapine for 30 mins before mounting in 1% LMA to reduce spontaneous motility. The Gcamp6f signal was recorded with 488nm laser intensity less than 0.5.

The zebrafish intestinal motility is quantified through recorded image series of zebrafish intestine using the method similar as previously described (Ganz et al., 2018). Intestinal µ velocity and ν velocity were used to estimate intestinal motility in zebrafish as previously described using the PIV-Lab MATLAB app (Ganz et al., 2018). A positive value of the µ velocity indicates an anterograde intestinal movement and a negative value of the µ velocity indicates a retrograde intestinal movement. The time-course µ velocity number is plotted as heatmaps. When calculating the mean velocity, only the mean velocity magnitude was calculated, which therefore doesn’t account for the movement direction. The MTrackJ FIJI plugin was used to quantify the mean velocity magnitude (Meijering et al., 2012).

To assess whether Trpa1^+^EEC activation induced intestinal motility change is due to the indirect communication through vagal afferent and efferent system, we anatomically disconnected zebrafish CNS with the intestine by decapitating. Optovin-treated unanesthetized zebrafish were mounted and placed on the 780 Zeiss upright confocal station as described above. The zebrafish head was then removed with a razor blade. The same imaging and 405nm activation of the mid-intestinal region was performed as described above.

### Enteric cholinergic neuron and vagal ganglion calcium imaging

*TgBAC(chata:Gal4); Tg(UAS:Gcamp6s); Tg(NBT:DsRed)* or *TgBAC(chata:Gal4); Tg(UAS:Gcamp6s); Tg(UAS:NTR-mCherry)* zebrafish were used to record *in vivo* calcium activity in enteric cholinergic neurons. The NBT promoter labels all ENS neurons while the Chata promoter labels only cholinergic enteric neurons. DsRed or mCherry fluorescence was used as reference for cholinergic neuron Gcamp quantification. Zebrafish larvae were incubated in 20 µM 4-DAMP for 30 mins before mounting in 1% LMA to reduce spontaneous motility and facilitate *in vivo* imaging using a Zeiss 780 upright confocal microscope with 20× water lenses. Serial images were taken at 5 s/frame. To record cholinergic neuron calcium activation, zebrafish was pretreated with Optovin and 40 nm light was applied at the frequency of 1 pulse/5s to the intestinal epithelium ROI. The Gcamp6s to DsRed fluorescence in cholinergic neurons was calculated for recorded.

*Tg(neurod1:Gcamp6f); Tg(neurod1:TagRFP)* zebrafish were used to record vagal sensory ganglia calcium activity *in vivo*. Zebrafish were anesthetized with 1 mg/mL α-Bungarotoxin (α-BTX) and gavaged with chemical compounds or bacteria as described (Naumann et al., 2016). Zebrafish larvae were mounted in 1% LMA and imaged under a Zeiss 780 upright confocal microscope. Z-stack images of the entire vagal ganglia were collected as serial images at 10 mins/frame and processed in FIJI. Individual vagal sensory neurons were identified and the Gcamp6f to TagRFP fluorescence ratios of individual vagal sensory neurons were calculated.

### Quantitative real-time PCR

Quantitative real-time PCR was performed as described previously (Murdoch et al., 2019). In brief, 20 zebrafish larvae digestive tracts were dissected and pooled into 1 mL TRIzol (ThermoFisher, 15596026). mRNA was then isolated with isopropanol precipitation and washed with 70% ethanol. 500ng mRNA was used for cDNA synthesis using the iScript kit (Bio-Rad, 1708891). Quantitative PCR was performed in triplicate 25 μL reactions using 2X SYBR Green SuperMix (PerfeCTa, Hi Rox, Quanta Biosciences, 95055) run on an ABI Step One Plus qPCR instrument using gene specific primers (Supplementary file 1). Data were analyzed with the ΔΔCt method. *18S* was used as a housekeeping gene to normalize gene expression.

### Mammalian TRPA1 activity analysis

HEK-293T cells were cultured in DMEM (Thermofisher Scientific, Waltham, MA) and supplemented with 10% fetal bovine serum (FBS) (Thermofisher Scientific), penicillin (100 units/mL) and streptomycin (0.1 mg/mL). Cells were plated on 100 mm tissue culture plates coated with poly-D-lysine (Sigma Aldrich, Saint Louis, MO) and grown to ∼60% confluence. The cells were transiently transfected for 16-24 hours with either human or mouse orthologs of TRPA1 using Fugene 6 transfection reagents and Opti-MEM (Thermofisher Scientific) according to the manufacturer’s protocol. Subsequently, cells were trypsinized, re-suspended and re-plated onto poly-D-lysine coated 96-well plates (Krystal black walled plates, Genesee Scientific) at 5×10^5^ cells/mL (100 µL/well) and allowed to grow for another 16-20 hrs prior to the experiments. Cells were maintained as monolayers in a 5% CO_2_ incubator at 37°C.

Measurements of changes in intracellular Ca^2+^ concentrations ([Ca^2+^]_i_) were performed as described previously (Caceres et al., 2017). In brief, cells in 96-well plates were loaded with Calcium 6, a no-wash fluorescent indicator, for 1.5 hrs (Molecular Devices, San Jose, CA) and then transferred to a FlexStation III benchtop scanning fluorometer chamber (Molecular Devices). Fluorescence measurements in the FlexStation were performed at 37°C (Ex:485 nm, Em: 525 nm at every 1.8 s). After recording baseline fluorescence, agonists (indole, IAld, cinnamaldehyde) were added and fluorescence was monitored for a total of 60 s. To determine the effects of TRPA1 inhibition on agonist response, TRPA1 transfected HEK-293 cells were pretreated with various concentrations of A967079 (Medchem101, Plymouth Meeting, PA), a specific antagonist of TRPA1, and then exposed to either 100 µM indole or IAld. The change in fluorescence was measured as Fmax-F0, where Fmax is the maximum fluorescence and F0 is the baseline fluorescence measured in each well. The EC50 and IC50 values and associated 95% confidence intervals for agonist (Indole and IAld) stimulation of Ca^2+^ influx and A967079 inhibition of agonist-induced Ca^2+^ influx, respectively, were determined by non-linear regression analysis with a 4-parameter logistic equation (Graphpad Prism, San Diego, CA). Indole and IAld concentration-response data was normalized to 1 mM cinnamaldehyde for EC50’s calculations and A967079 concentration-response data was normalized to 100 µM indole or IAld for IC50’s calculations.

### HPLC-MS analysis of Trp-Indole derivatives

The chemical profiling of Trp-Indole derivatives was performed using 1 L culture of *E. tarda*. The strain was inoculated in 3 mL of TSB medium and cultivated for 1 day on a rotary shaker at 180 rpm at 30°C under aerobic conditions. After 1 day, 1 mL of E. tarda liquid culture was inoculated in 1 L of TSB medium in a 4-L Pyrex flask. The *E. tarda* culture was incubated at 30°C for 24 hr under aerobic conditions. For time-course screening, 10 mL from the *E. tarda* TSB culture was collected at 0, 6, 18, and 24 hours. Each 10 mL sample of *E. tarda* culture was extracted with 15 mL of ethyl acetate (EtOAc). The EtOAc layer was separated from the aqueous layer and residual water was removed by addition of anhydrous sodium sulfate. Each EtOAc fraction was dried under reduced pressure, then resuspended in 500 μL of 50% MeOH/50% H_2_O and 50 μL of each sample were analyzed using an Agilent Technologies 6130 quadrupole mass spectrometer coupled with an Agilent Technologies 1200–series HPLC (Agilent Technologies, Waldbron, Germany). The chemical screening was performed with a Kinetex® EVO C18 column (100 × 4.6 mm, 5 µm) using the gradient solvent system (10 % ACN/90 % H_2_O to 100 % ACN over 20 min at a flow rate of 0.7 mL/min).

For HPLC-MS analysis of *E. tarda* in GZM medium, the remaining 1 L culture of *E. tarda* in TSB culture was centrifuged at 7,000 rpm for 30 min. Pellets were transferred to 1 L of GZM medium in a 4-L Pyrex flask and cultivated on a rotary shaker at 30°C for 24 hr. For time-course screening, 10 mL from the *E. tarda* GZM culture was collected at 0, 1, 6, and 24 hours. Sample preparation and HPLC-MS analysis of *E. tarda* culture GZM medium were performed using same procedures as described above for TSB. Trp-Indole derivatives of *E. tarda* culture broths were identified by comparing the retention time and extracted ion chromatogram with authentic standards. Extracted ions were selected for Indole (m/z 117, Sigma-Aldrich), IAld (m/z 145, Sigma-Aldrich), IAAld (m/z 159, Ambeed), IEt (m/z 161, Sigma-Aldrich), IAM (m/z 174, Sigma-Aldrich), IAA (m/z 175, Sigma-Aldrich), and IpyA (m/z 203, Sigma-Aldrich).

For HPLC-MS analysis of Trp-indole derivatives from 15 different bacterial strains in TSB medium, each of the strains (*Acinetobacter* sp. ZOR0008, *Aeromonas veronii* ZOR0002, *Bacillus subtilis* 168, *Chryseobacterium* sp. ZOR0023, *Edwardsiella tarda* 15974, *Edwardsiella tarda* 23685, *Edwardsiella tarda* LSE40, *Edwardsiella* tarda FLG6-60, *Enterobacter* sp. ZOR0014, *Escherichia coli* MG1655, *Exiguobacterium acetylicum* sp. ZWU0009, *Plesiomonas* sp. ZOR0011, *Pseudomonas aeruginosa* PAK, *Shewanella* sp. ZOR0012, and *Vibrio* sp. ZWU0020) were inoculated in 3 mL of TSB medium and cultivated for 1 day on a rotary shaker at 180 rpm at 30°C under aerobic conditions. After 1 day, 1 mL of each liquid culture was inoculated in 100 mL of TSB medium in 500 mL Pyrex flasks and cultivated on a rotary shaker at 30°C overnight. A 10 mL sample was taken from each culture and extracted and analyzed via HPLC-MS as explained above. CFU was calculated for each bacterial liquid culture and the HPLC-MS data was normalized to the CFU.

For HPLC-MS analysis of Trp-indole derivatives from 15 different bacterial strains in GZM medium, the remaining 100 mL culture of each strain was centrifuged at 4500 rpm for 20 min. Pellets were transferred to 100 mL of GZM medium in 500 mL Pyrex flasks and cultivated on a rotary shaker at 30°C overnight. Sample preparation and HPLC-MS analysis of each GZM culture were performed using the same procedure as described above.

For HPLC-MS analysis of Trp-indole derivatives from murine small intestine and large intestine, three 10-week old female and three 10-week old male conventionally-reared specific pathogen-free C57BL/6J mice were ordered from Jackson Lab. The mice were not fast in advance and euthanized with 5% isoflurane. The 2/5-4/5 portion of the small intestinal region and the colon caudal to cecum was collected from each mouse and transferred to a 50 mL conical tube that was placed on dry-ice. 80% methanol was then added according to the tissue weight (50 µL/mg tissue). The intestine was then homogenized with a Tissue-Tearor (BioSpec Products, 985370). Following homogenizing, the tryptophan metabolites were extracted and analyzed with HPLC-MS as explained above. The relative metabolite abundance was normalized to tissue weight. These mouse experiments conformed to the US Public Health Service Policy on Humane Care and Use of Laboratory Animals, using protocol number A170-17-07 approved by the Institutional Animal Care and Use Committee of Duke University.

### Measurement of serotonin release from mouse and human small intestine

These experiments using C57BL/6J mice were approved by the Flinders University Animal Welfare Committee (number 965-19) and human ileum tissue was collected from resected small and large intestine from patients that gave written informed consent under the approval of the Southern Adelaide Clinical Human Research Ethics Committee (number 50.07) as previous (Sun et al., 2019a). Mice were euthanized at 8 to 12 weeks by isoflurane overdose followed by cervical dislocation. The duodenum was removed and placed in Krebs solution oxygenated with 95 % O_2_, 5 % CO_2_. A midline incision was made along the duodenum to create a flat sheet, the section was pinned mucosal-side up in an organ bath lined with Sylgard and containing oxygenated Krebs solution. Serotonin release was measured using amperometry. A carbon-fibre electrode (5-μm diameter, ProCFE; Dagan Corporation, Minneapolis, MN), was lowered above the mucosa and 400 mV potential was applied to the electrode causing oxidation of serotonin (Zelkas et al., 2015). 10mM Indole and/or 50µM HC030031 were applied to tissue by constantly perfusing the bath. The change in amplitude due to serotonin oxidation was recorded using an EPC-10 amplifier and Pulse software (HEKA Electronic, Lambrecht/Pfalz, Germany), and samples at 10 kHz and low-pass filtered at 1 kHz. Data was assessed as peak current during each treatment. Data was analyzed comparing all groups using one-way ANOVA with Tukey’s post-hoc test. For the mouse experiments, 6 independent experiments were performed in 6 mouse duodenal samples. For the human experiments, 4 independent experiments were performed in 3 human samples.

### Statistical analysis

The appropriate sample size for each experiment was suggested by preliminary experiments evaluating variance and effects. Using significance level of 0.05 and power of 90%, a biological replicate sample number 8 was suggested for EEC CaMPARI analysis. For each experiment, wildtype or indicated transgenic zebrafish embryos were randomly allocated to test groups prior to treatment. Individual data points, mean and standard deviation are plotted in each figure. The raw data points in each figure are represented as solid dots. Data were analyzed using GraphPad Prism 7 software. For experiments comparing just two differentially treated populations, a Student’s t-test with equal variance assumptions was used. For experiments measuring a single variable with multiple treatment groups, a single factor ANOVA with post hoc means testing (Tukey) was utilized. Statistical evaluation for each figure was marked * P<0.05, ** P<0.01, *** P<0.001, **** P<0.0001 or ns (no significant difference, P>0.05).

**SUPPLEMENTAL TABLES AND VIDEOS**

**Table S1. Zebrafish EEC RNA-seq data analyzed by DEseq2, related to Figure 2**.

**Table S2. Comparison of zebrafish EECs with human and mouse EECs using RNA-seq, related to Figure 2**.

**Table S3. Expression of hormones, transcription factors, receptors, and innate immune genes in EECs and other IECs, related to Figure 2**.

**Video 1. *E. tarda* activates EECs *in vivo*, related to Figure 1**. Time-course video of *Tg(neurod1:Gcamp6f)* zebrafish stimulated with *E. tarda* bacteria. Anterior is to the right, and dorsal is to the top.

**Video 2. Trpa1 agonist activates EECs *in vivo*, related to Figure 2**. Time-course videos of *trpa1b*+/+ and *trpa1b*-/-*Tg(neurod1:Gcamp6f)* zebrafish stimulated with Trpa1 agonist AITC. Anterior is to the right, and dorsal is to the top.

**Video 3. Optovin-UV activates EECs, related to Figure 4**. Time-course video of *Tg(neurod1:Gcamp6f); Tg(neurod1:TagRFP)* zebrafish before and post Optovin-UV induced EEC Trpa1 activation. Anterior is to the left, and dorsal is to the bottom.

**Video 4. Activation of EEC Trpa1 in control zebrafish increases intestinal motility, related to Figure 4**. Time-course video of *Tg(neurod1:Gcamp6f)* WT zebrafish before and post Optovin-UV induced EEC Trpa1 activation. Anterior is to the left, and dorsal is to the top.

**Video 5. Activation of EEC Trpa1 in EEC ablated zebrafish does not increase intestinal motility, related to Figure 4**. Time-course video of *Tg(neurod1:Gcamp6f)* EEC ablated zebrafish before and post Optovin-UV induced EEC Trpa1 activation. Anterior is to the left, and dorsal is to the top.

**Video 6. Optic EEC activation in EEC-ChR2 expressing transgenic zebrafish, related to Figure 4**. Time-course video of *Tg(neurod1:Gcamp6f); Tg(neurod1:Gal4); Tg(UAS:ChR2-mCherry)* zebrafish before and post yellow light induced EEC activation. Anterior is to the left, and dorsal is to the bottom.

**Video 7. Activation of Trpa1+ChR2+EECs increases intestine motility, related to Figure 4**. Time-course videos of *TgBAC(trpa1b:EGFP); Tg(neurod1:Gal4); Tg(UAS:ChR2-mCherry)* before and post optic activation of Trpa1-EECs and Trpa1+ EECs. Anterior is to the left, and dorsal is to the bottom. The yellow light was delivered specifically to the selected EECs to activate the ChR2 channel. Note that the first frame in each video shows the EGFP channel to identify EECs that do or do not express *trpa1b*. Also note that the ChR2 EECs are in the intestinal bulb and an anterograde intestinal movement was observed upon Trpa1+ChR2+EEC activation.

**Video 8. Activation of middle intestinal Trpa1+ChR2+EECs increases intestine motility, related to Figure 4**. Time-course videos of *TgBAC(trpa1b:EGFP); Tg(neurod1:Gal4); Tg(UAS:ChR2-mCherry)* before and post optic activation of Trpa1+ EECs. Anterior to the left. Note that the Trpa1+ChR2+EECs are in the posterior intestine and activation of the posterior intestinal Trpa1+ChR2+EECs induces anterograde intestinal movement.

**Video 9. *E. tarda* increases intestinal motility, related to Figure 4**. Time-course videos of WT zebrafish 30 mins post *Aeromonas* sp. or *E. tarda* gavage. Anterior is to the left, and dorsal is to the top.

**Video 10. EECs physically connect to Chata+ enteric neurons, related to Figure 5**. 3D-reconstruction of *Tg(chata: NTR-mCherry)* zebrafish intestine stained with 2F11 antibody that labels EEC. The Chata+ nerve is shown as magenta and the EECs are shown as green.

**Video 11. Activation of EEC Trpa1 increases Chata+ ENS calcium, related to Figure 5**. Time-course videos of *TgBAC(chata:Gal4); Tg(UAS;Gcamp6s)* that before and post Optovin-UV induced EEC Trpa1 activation. Anterior is to the left, and dorsal is to the top.

**Video 12. *E. tarda* increases vagal ganglia calcium, related to Figure 6**. Time-course videos of vagal ganglia calcium in *Tg(neurod1:Gcamp6f); Tg(neurod1:TagRFP)* zebrafish that are gavaged PBS or *E. tarda*. Anterior is to the left, and dorsal is to the top.

**Video 13. IAld activates EECs *in vivo*, related to Figure 7**. Time-course videos of *trpa1b*+/+ (WT) and *trpa1b*-/-*Tg(neurod1:Gcamp6f)* zebrafish that are stimulated with IAld. Note that there is a basal amount of intestinal motility associated with this methylcellulose preparation that is retained in vehicle-only negative controls (not shown) and in trpa1b mutants. Anterior is to the right, and dorsal is to the top.

**Video 14. Indole activates EECs *in vivo*, related to Figure 7**. Time-course videos of *trpa1b*+/+ (WT) and *trpa1b*-/-*Tg(neurod1:Gcamp6f)* zebrafish that are stimulated with indole. Note that there is a basal amount of intestinal motility associated with this methylcellulose preparation that is retained in vehicle-only negative controls (not shown) and in trpa1b mutants. Anterior is to the right, and dorsal is to the top.

## SUPPLEMENTAL FIGURES

**Figure S1.**
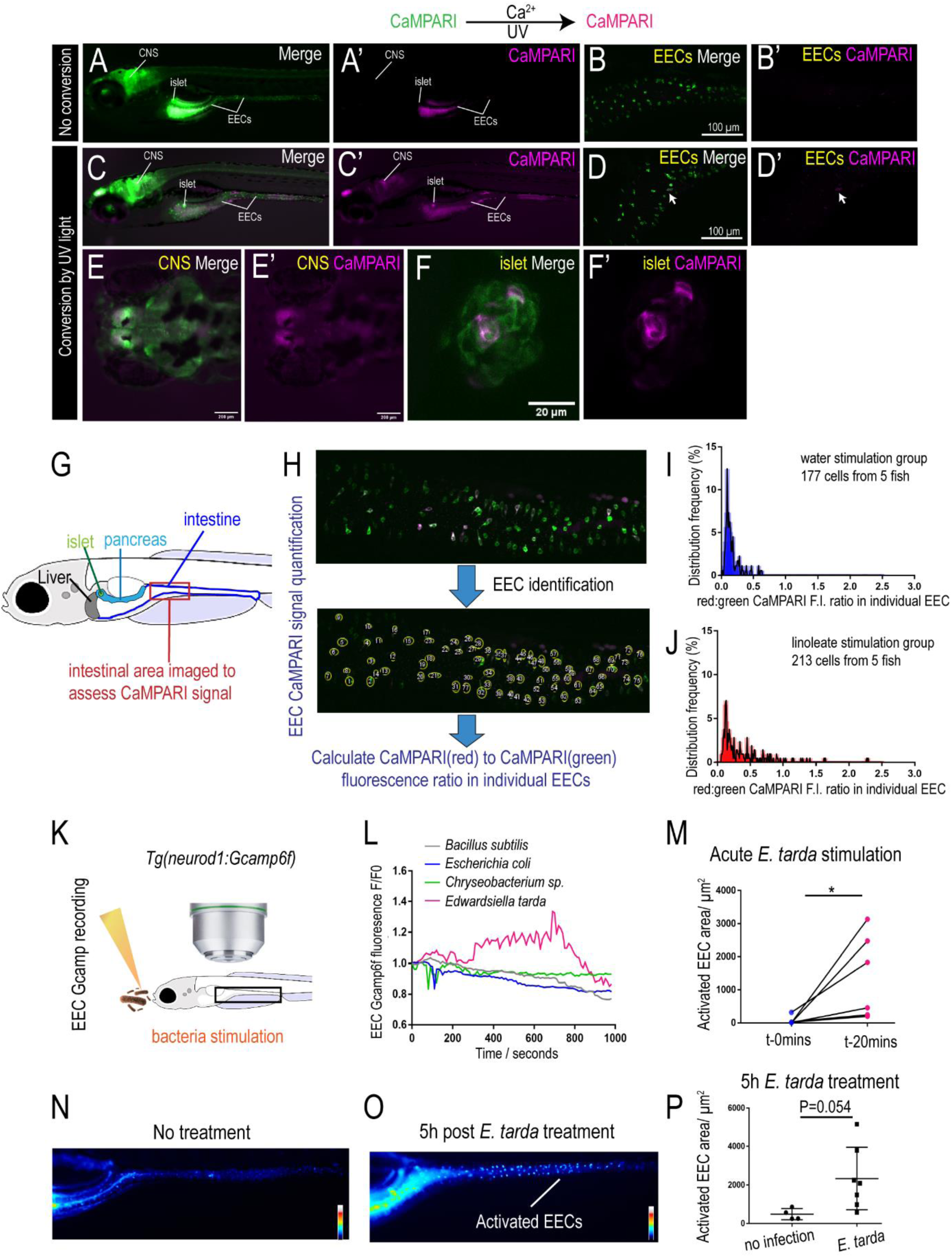
*E. tarda* activates EECs *in vivo*, related to Main Figure 1. (A) Epifluorescence image of *Tg(neurod1:CaMPARI)* zebrafish without UV conversion. Note that there is no red CaMPARI signal (magenta) in A’. (B) Confocal image of intestinal EECs in *Tg(neurod1:CaMPARI)* zebrafish without UV conversion. (C) Epifluorescence image of unstimulated *Tg(neurod1:CaMPARI)* zebrafish post UV conversion. The red CaMPARI signal is apparent in CNS and islets in C’. (D-F’) Confocal image of intestinal EECs (D, D’), CNS (E, E’) and islets (F, F’) in unstimulated *Tg(neurod1:CaMPARI)* zebrafish after UV conversion. (G) Schematic of liver, pancreas and intestine in 6 dpf zebrafish larvae. The intestinal region that is imaged to assess the CaMPARI signal is indicated by a red box. (H-J) Quantification of EEC red:green CaMPARI fluorescence ratio in water- and linoleate-stimulated zebrafish. (K) Schematic of *in vivo* EEC Gcamp recording in response to bacterial stimulation in *Tg(neurod1:Gcamp6f)* zebrafish. (L) Quantification of EEC Gcamp6f fluorescence in response to stimulation by different bacteria. (M) Quantification of EEC Gcamp6f fluorescence before and 20 mins after *E. tarda* administration. (N-O) Fluorescence image of zebrafish intestine in *Tg(neurod1:Gcamp6f)* zebrafish without treatment (N) or 5 hours post *E. tarda* treatment (O). (P) Quantification of EEC Gcamp6f fluorescence in zebrafish without or with *E. tarda* treatment. Student’s t-test was used in M and P for statistical analysis. * p<0.05.

**Figure S2.**
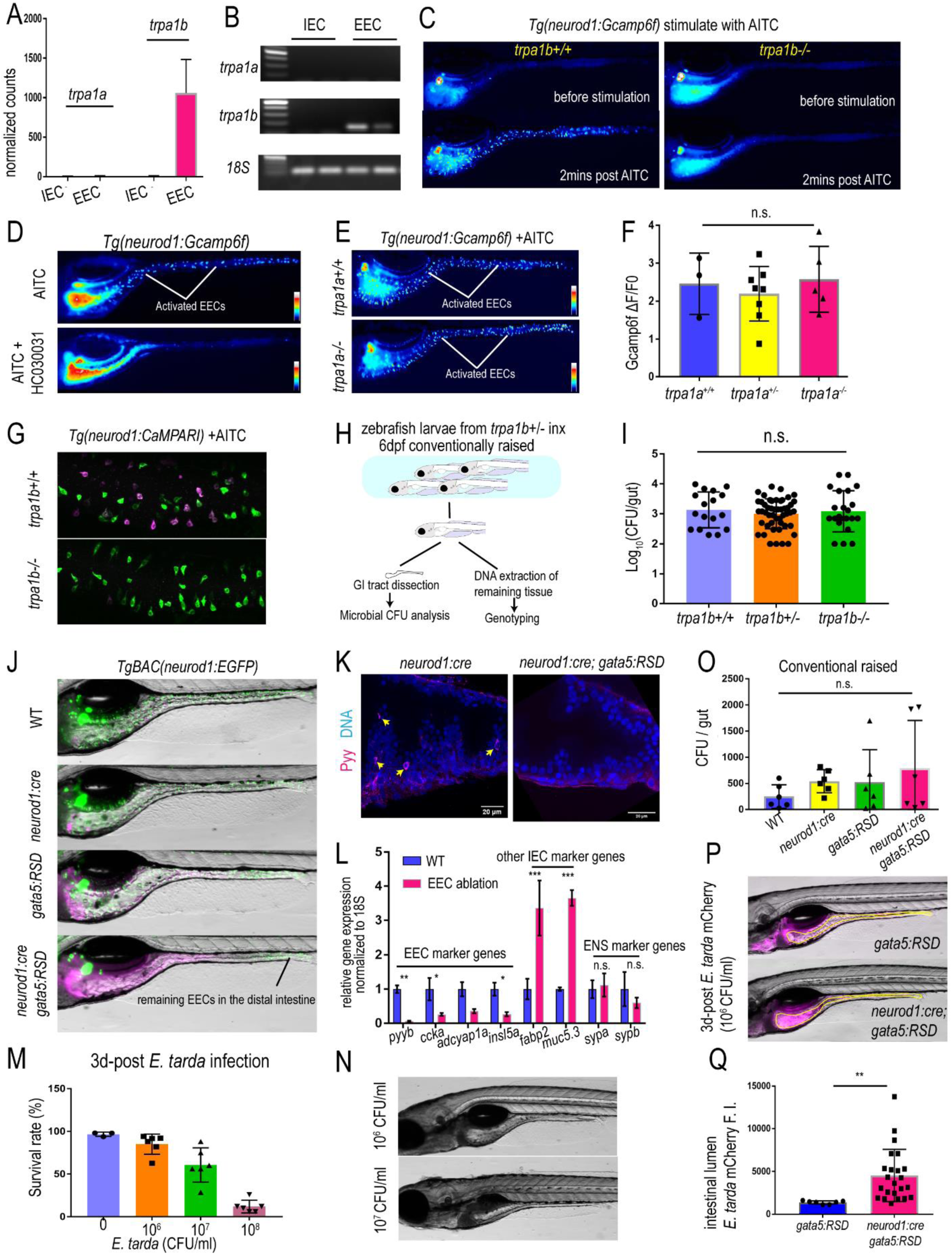
EECs express *trpa1b* and respond to Trpa1 agonist, related to Main Figure 2 and Figure 3. (A) Normalized counts of *trpa1a* and *trpa1b* gene expression in zebrafish EECs and other IECs from zebrafish EEC RNA-seq data (Table S1). (B) Gel image of PCR product from FACS sorted EECs and other IECs cell population using primers from *trpa1a, trpa1b* and *18S*. (C) Epifluorescence image of *trpa1b*+/+ (left) and *trpa1b*-/-(right) *Tg(neurod1:Gcamp6f)* zebrafish before or 2 mins post Trpa1 agonist AITC stimulation. (D) Epifluorescence image of *Tg(neurod1:Gcamp6f)* zebrafish following AITC stimulation with or without Trpa1 antagonist HC030031 treatment. (E) Epifluorescence image of *trpa1a*+/+ and *trpa1a*-/-*Tg(neurod1:Gcamp6f)* zebrafish 2 mins after AITC stimulation. (F) Quantification of EEC Gcamp fluorescence signal in *trpa1a*+/+, *trpa1a*+/- and trpa1a-/-zebrafish. (G) Confocal projection of *trpa1b*+/+ and *trpa1b*-/-*Tg(neurod1:CaMPARI)* zebrafish after AITC stimulation and UV light photoconversion. (H) Model of gut bacterial CFU quantification. (I) Quantification of gut bacterial CFU in *trpa1b*+/+, *trpa1b*+/- and *trpa1b*-/-conventionalized zebrafish. (J) Epifluorescence image of WT, *Tg(neurod1:cre), Tg(gata5:RSD)* and *Tg(neurod1:cre); Tg(gata5:RSD)* zebrafish. The EECs in all the groups are labelled by *Tg(neurod1:EGFP)*. Note that *neurod1:EGFP* labelling is largely absent in *Tg(neurod1:cre); Tg(gata5:RSD)* zebrafish indicating EEC ablation. (K) Confocal images of *Tg(neurod1:cre)* (left) and *Tg(neurod1:cre); Tg(gata5:RSD)* (right) zebrafish intestine stained with PYY antibody. Yellow arrows in D indicate PYY+ EECs. (L) qPCR analysis of EEC marker genes, other IEC marker genes and neuronal genes in WT and EEC-ablated zebrafish. (M) Quantification of zebrafish survival rate when treated with different doses of *E. tarda* FL6-60. (N) Representative image of zebrafish treated with 10^6^ CFU/ml or 10^7^ CFU/ml *E. tarda*. Note that in the 10^6^ CFU/ml treated zebrafish, the majority of the surviving zebrafish do not exhibit gross pathology (top image). While many of the survived zebrafish treated with 10^7^ CFU/ml *E. tarda* displayed deflated swim bladder, altered intestinal morphology and ruptured skin (bottom image). (O) Quantification of gut bacterial CFU in WT, *Tg(neurod1:cre), Tg(gata5:RSD)* and *Tg(neurod1:cre); Tg(gata5:RSD)* conventionalized zebrafish. (P) Epifluorescence image of *Tg(gata5:RSD)* and EEC-ablated zebrafish treated with *E. tarda* mCherry for 3 days. (Q) Quantification of *E. tarda* mCherry fluorescence intensity in the intestinal lumen of Tg*(gata5:RSD)* or EEC-ablated zebrafish. One-way ANOVA with Tukey’s post test was used in F, I, L, O and student t-test was used in Q for statistical analysis. n.s. (not significant), *p<0.05, **p<0.01, ***p<0.001.

**Figure S3.**
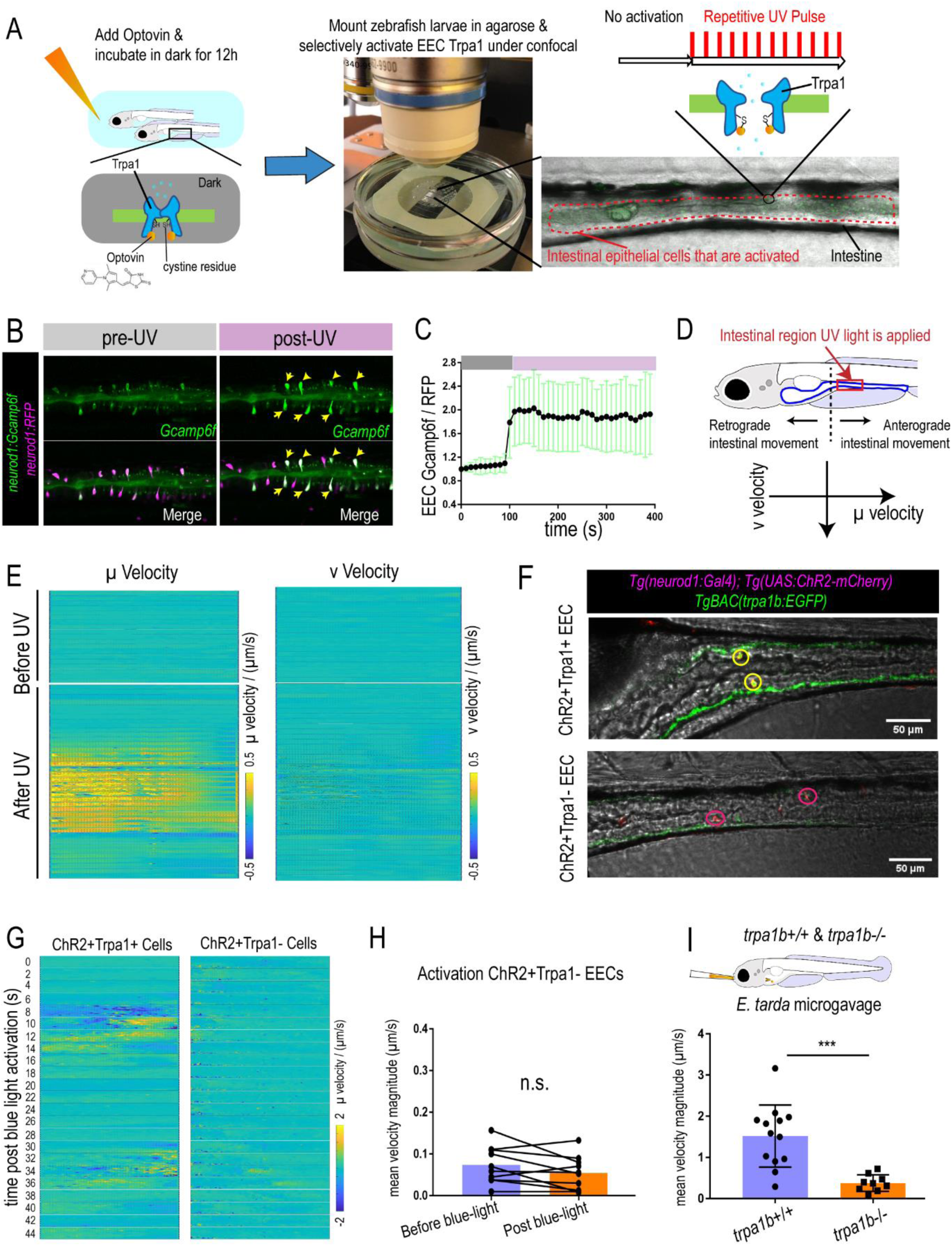
Activation of EEC Trpa1 signaling promotes intestinal motility, related to Main Figure 4. (A) Experimental design for activating EEC Trpa1 signaling using Optovin-UV. (B) Confocal image of *Tg(neurod1:Gcamp6f); Tg(neurod1:TagRFP)* zebrafish intestine before (images on the left) and after (images on the right) UV light activation. Yellow arrows indicate the subpopulation of EECs exhibiting increased Gcamp fluorescence following UV activation. (C) Quantification of the EEC Gcamp6f to TagRFP fluorescence ratio before and after UV activation. (D) Schematic of intestinal movement in larval zebrafish. The proximal zebrafish intestine exhibits retrograde movement while mid-intestine and distal intestine exhibit anterograde movement. The imaged and UV light activated intestinal region in the Optovin-UV experiment is indicated by the red box. The µ velocity indicates intestinal horizontal movement. A positive value indicates anterograde movement and a negative value indicates retrograde movement. The ν velocity indicates intestinal vertical movement. (E) Quantification of intestinal motility using PIV-LAB velocity analysis before and after UV activation. Note that Optovin-UV induced Trpa1 activation increased µ velocity (horizontal movement) more than ν velocity (vertical movement). (F) Confocal image of ChR2+Trpa1+ EECs (yellow circles, top image) and ChR2+Trpa1-EECs (red circles, bottom image) in *TgBAC(trpa1b:EGFP); Tg(neurod1:Gal4); Tg(UAS:ChR2-mCherry)* zebrafish. (G) Quantification of µ velocity following blue light activation of ChR2+Trpa1+ or ChR2+Trpa1-EECs. (H) Quantification of mean intestinal velocity magnitude change before and after blue light activation of ChR2+Trpa1-EECs. (I) Quantification of mean intestinal velocity magnitude in response to *E. tarda* gavage in *trpa1b*+/+ or *trpa1b*-/-zebrafish. Student t-Test was used in I. *** P<0.001.

**Figure S4.**
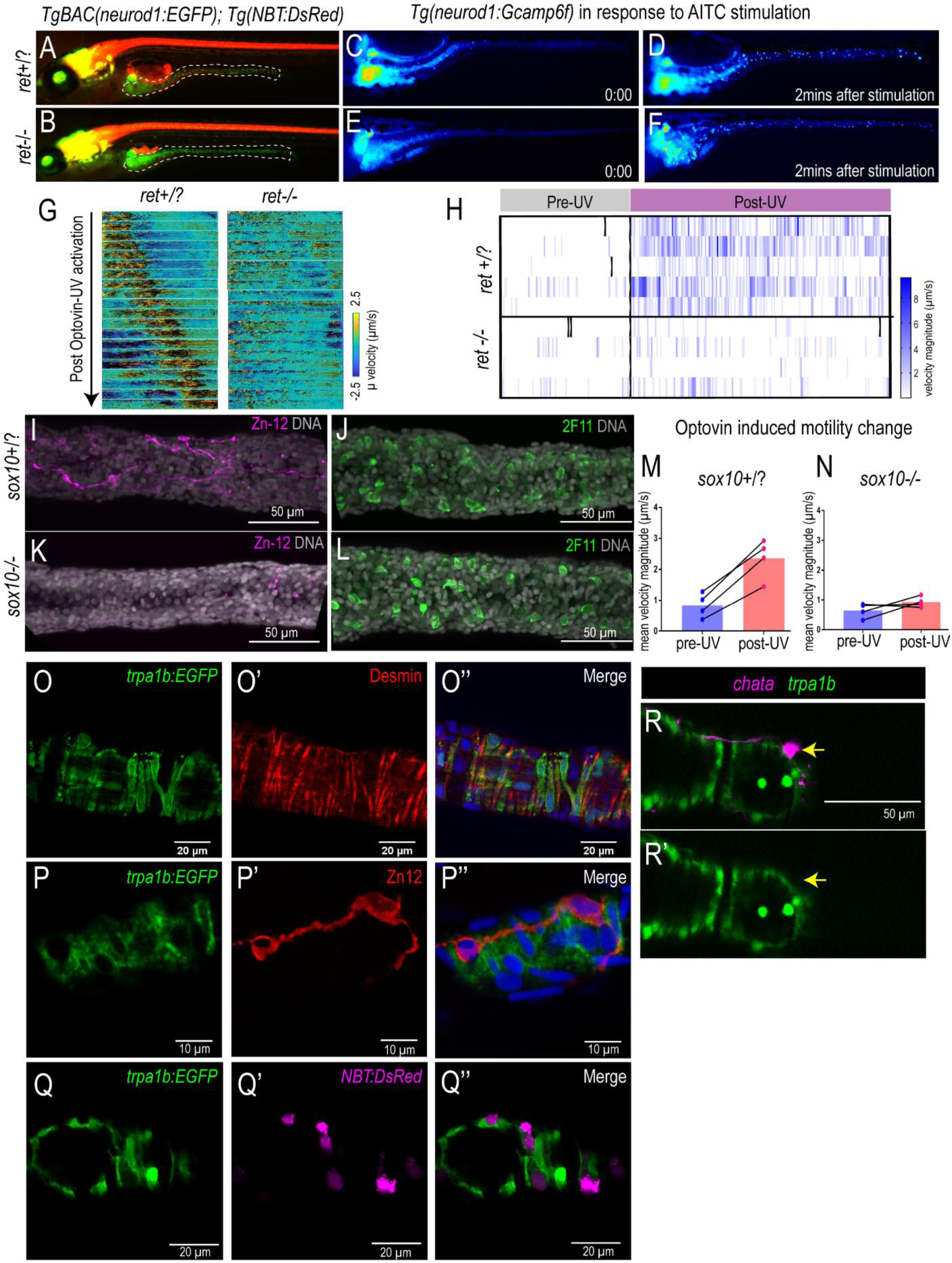
The role of the enteric nervous system in EEC Trpa1-induced intestinal motility, related to Main Figure 5. (A-B) Epifluorescence image of *ret+/+ or ret+/-* (*ret*+/?, A) and *ret*-/-(B) *Tg(NBT:DsRed); Tg(neurod1:EGFP)* zebrafish. The intestines are denoted by white dash lines. (C-D) Epifluorescence image of *ret*+/? *Tg(neurod1:Gcamp6f)* zebrafish before (C) and 2 mins after AITC stimulation (D). (E-F) Epifluorescence image of *ret*-/-*Tg(neurod1:Gcamp6f)* zebrafish before (E) and 2 mins after AITC stimulation (F). (G) Quantification of *ret*+/? and *ret*-/-intestinal µ velocity following Optovin-UV-induced Trpa1 activation. (H) Quantification of velocity before and after Optovin-UV-induced Trpa1 activation in *ret*+/? and *ret*-/-zebrafish. (I-J) Confocal projection of *sox10*+/? zebrafish intestine stained with Zn12 (I, magenta, ENS labeling) or 2F11(J, green, EEC labeling). (K-L) Confocal projection of *sox10*-/-zebrafish intestine stained with zn-12 (K) or 2F11(L). (M-N) Quantification of changes in mean intestinal velocity magnitude before and after Optovin-UV activation in *sox10*+/? (M) or *sox10*-/-(N) zebrafish. (O-P) Confocal projection of *TgBAC(trpa1b:EGFP)* zebrafish intestine stained with Desmin (myoblast or smooth muscle cell marker, O’) or Zn12 (ENS marker, P’). (Q) Confocal image of *TgBAC(trpa1b:EGFP); Tg(NBT:DsRed)* zebrafish intestine. Note in P and Q that both Zn12+ ENS and NBT+ ENS are *trpa1b*-. (R) Confocal image of *TgBAC(chata:Gal4); Tg(UAS:NTR-mCherry); TgBAC(trpa1b:EGFP)* zebrafish intestine. Note that the *chata*+ ENS are *trpa1b*-.

**Figure S5.**
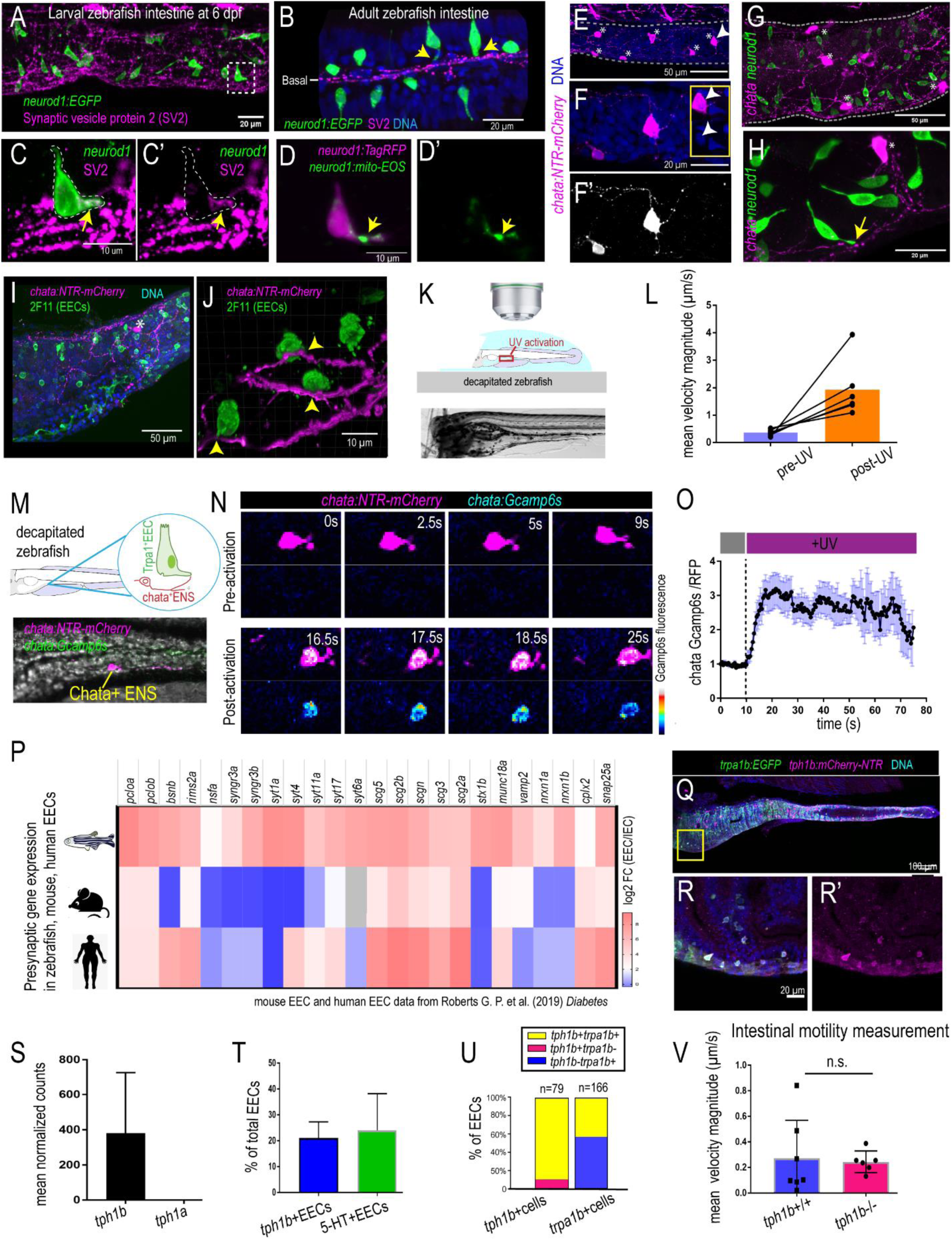
Zebrafish EECs directly communicate with *chata*+ ENS, related to Main Figure 5. (A-B) Confocal projection of 6 dpf (A) and adult (B) *Tg(neurod1:EGFP)* zebrafish intestine stained with the neuronal marker synaptic vesical protein 2 (SV2, magenta) antibody. (C) Higher magnification view of an EEC that exhibiting a neuropod contacting SV2 labelled neurons in the intestine. Yellow arrow indicates the EEC neuropod is enriched in SV2. (D) Higher magnification view of an EEC and neuropod in *Tg(neurod1:TagRFP); Tg(neurod1:mitoEOS)* zebrafish. The yellow arrow indicates the EEC neuropod is enriched in mitochondria (green, labelled by *neurod1:mitoEOS*). (E) Confocal projection of *chata*+ ENS in *TgBAC(chata:Gal4); Tg(UAS:mCherry-NTR)* zebrafish intestine. Asterisks indicate the *chata*+ enteric neuron cell bodies. (F) Higher magnification view of a *chata*+ ENS (white arrow in E). The nuclei of this *chata*+ enteric neuron is shown on the right. (F’) The axon processes of the *chata*+ enteric neuron. Note this neuron displays a typical Dogiel type II morphology in which multiple axons project from the cell body. (G) Confocal projection of *chata*+ ENS and EECs in *TgBAC(chata:Gal4); Tg(UAS:mCherry-NTR); Tg(neurod1:EGFP)* zebrafish intestine. EECs are labeled as green and *chata*+ ENS are labeled as magenta. Asterisks indicate the *chata*+ enteric neuron cell bodies. (H) Higher magnification view of the physical connection between EECs and the *chata*+ enteric neuron. Yellow arrow indicates an EEC forming a neuropod to contact a *chata*+ enteric neuron. (I) Confocal projection of EECs (2F11+, green) and *chata*+ ENS (magenta) in *TgBAC(chata:Gal4); Tg(UAS: NTR-mCherry)* zebrafish. An asterisk indicates the *chata*+ ENS cell body. (J) Higher magnification view of the connection between EECs and *chata*+ ENS fibers. The point where EECs connected with *chata*+ nerve fibers are indicated by yellow arrows. (K) Schematic of Optovin-UV experiment in zebrafish that are anatomically disconnected from their CNS. The Optovin-treated zebrafish were mounted and placed on a confocal objective station. Immediately prior to imaging, the head of the mounted zebrafish was quickly removed with a sharp razor blade and imaging was then performed. (L) Quantification of the mean intestinal velocity before and post UV treatment in decapitated zebrafish. (M) Schematic and confocal image shows the *chata*+ ENS which is labelled by both *Gcamp6s* and *mCherry* in decapitated *TgBAC(chata:Gcamp6s); Tg(UAS:Gcamp6s); Tg(UAS:NTR-mCherry)* zebrafish. (N) Confocal image shows the *chata*+ ENS Gcamp fluorescence intensity before and post Trpa1^+^ EEC activation by UV light. (O) Quantification of relative *chata*+ ENS Gcamp fluorescence intensity before and post Trpa1+ EEC activation. The Gcamp fluorescence intensity was normalized to mCherry fluorescence. n=12 from 7 zebrafish. (P) Log_2_ fold change of presynaptic genes in zebrafish, mouse and human EECs (Table S2). (Q-R) Confocal image and higher magnification view of *TgBAC(trpa1b:EGFP); Tg(tph1b:mCherry-NTR)* zebrafish intestine showing the *tph1b*+ (magenta) *trpa1b*+ (green). (S) Quantification of *tph1a* and *tph1b* in zebrafish EECs. (T) Quantification of 5-HT+ or *tph1b*+ EECs. (U) Quantification of *tph1b*+ and *trpa1b*+ EECs. Note the majority of *tph1b*+ EECs are *trpa1b*+. (V) Quantification of mean intestinal µ velocity in unstimulated *tph1b*+/- and *tph1b*-/-zebrafish. Student t-test was used in V.

**Figure S6.**
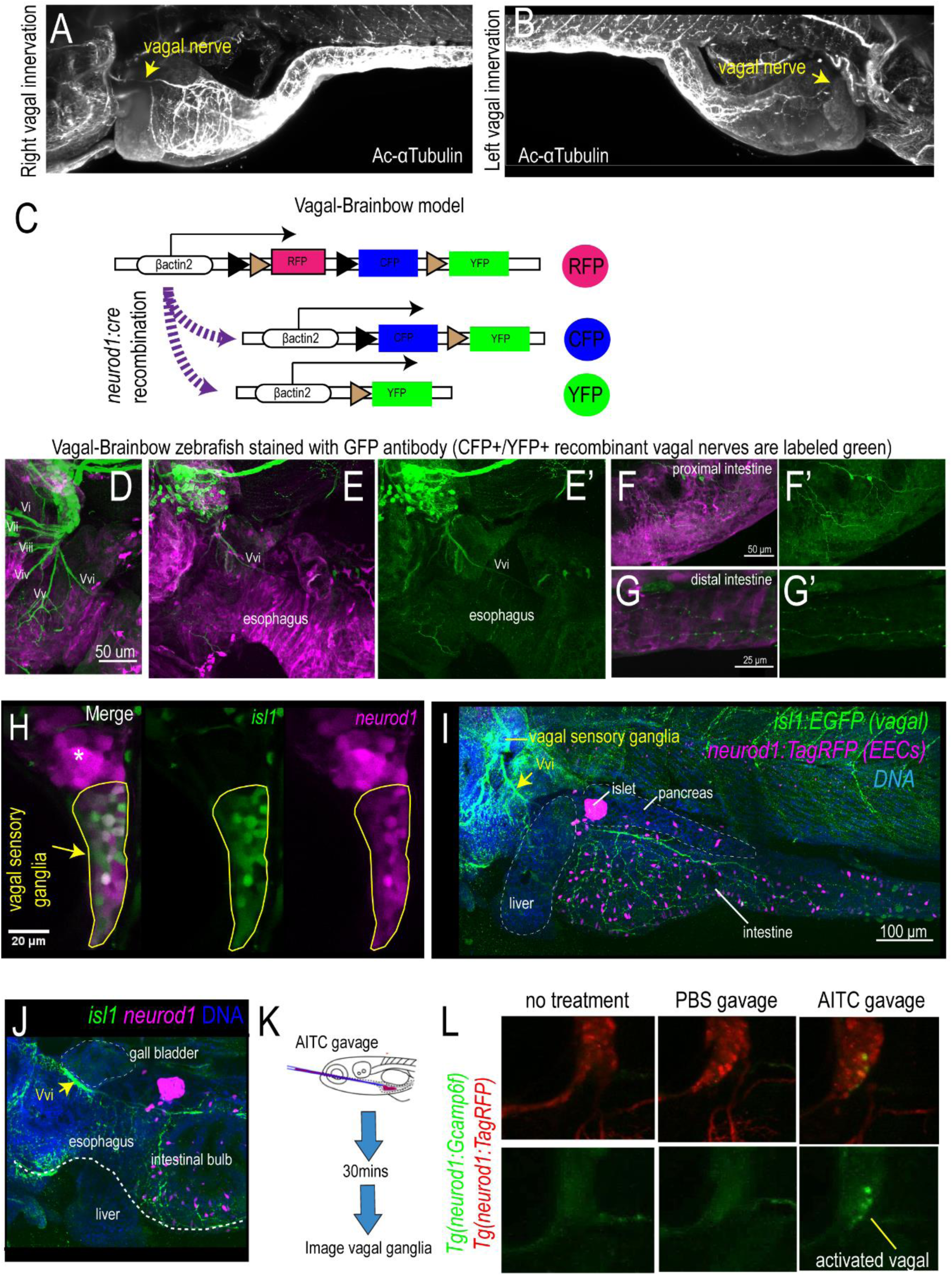
Zebrafish vagal sensory nerve innervate the intestine, related to Main Figure 6. (A-B) Lightsheet imaging of the right (A) and left (B) side of zebrafish intestine stained with acetylated α-tubulin antibody (white). (C) Schematic diagram of the Vagal-Brainbow model to label vagal sensory cells using *Tg(neurod1:cre); Tg(βact2:Brainbow)* zebrafish. See Vagal-Brainbow projection in Fig. 6F. (D) Confocal image of vagal ganglia in brainbow zebrafish stained with GFP antibody (green). Note that GFP antibody recognizes both YFP+ and CFP+ vagal sensory neurons. Six branches (V_i_ to V_vi_) extend from the vagal sensory ganglia and branch V_vi_ innervates the intestine. (E-E’) Confocal image of vagal sensory ganglia in brainbow zebrafish showing that V_vi_ exits from the ganglia and courses behind the esophagus. (F-G) Confocal image of the proximal (F) and distal (G) intestine in brainbow zebrafish. The vagus nerve (green) innervates both intestinal regions. (H) Confocal image of vagal sensory ganglia in *Tg(isl1:EGFP); Tg(neurod1:TagRFP)* zebrafish. The vagal sensory ganglia is indicated by a yellow circle. The asterisk indicates the posterior lateral line ganglion. Note that *isl1* (green) is expressed in the vagal sensory ganglia and overlaps with *neurod1* (magenta). (I) Confocal image of intestine in *Tg(isl1:EGFP); Tg(neurod1:TagRFP)* zebrafish. The vagus nerve is labelled by *isl1* (green) and the intestinal EECs are labelled by *neurod1* (magenta). (J) Confocal plane of intestine in *Tg(isl1:EGFP); Tg(neurod1:TagRFP)* zebrafish. Note that the V_vi_ branch of the vagus nerve is labelled by *isl1* and travels behind the esophagus to innervate the intestine. (K) Schematic of *in vivo* vagal calcium imaging in PBS or AITC gavaged zebrafish. (L) *In vivo* vagal calcium imaging of *Tg(neurod1:TagRFP); Tg(neurod1:Gcamp6f)* zebrafish without gavage, gavaged with PBS or gavaged with AITC.

**Figure S7.**
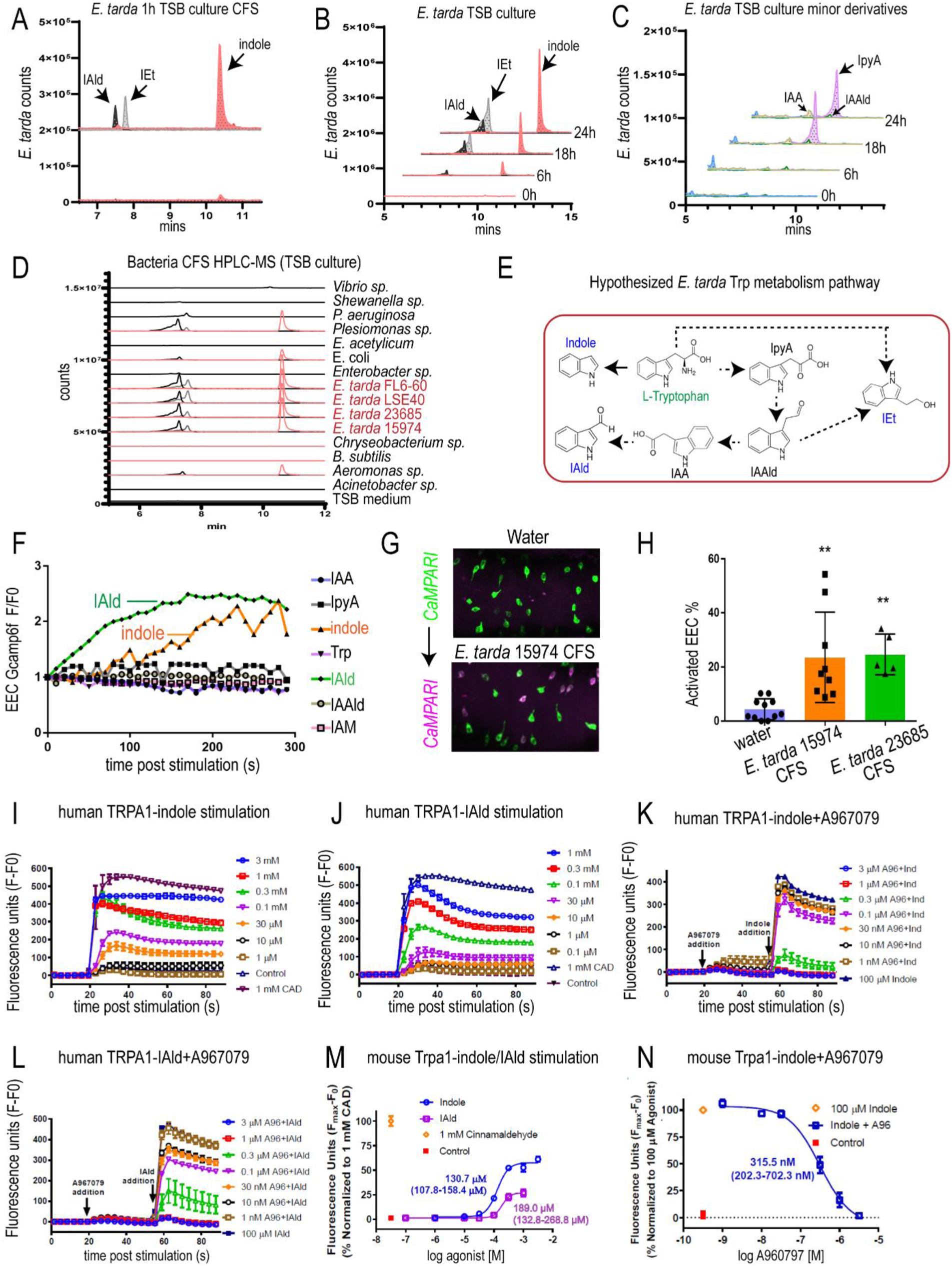
*E. tarda* secretes tryptophan catabolites indole and IAld that activate Trpa1, related to Main Figure 7. (A) Chemical profiles of Trp-Indole derivatives from supernatants of *E. tarda* in nutrient-rich TSB media. (B) Screening of supernants of *E. tarda* in TSB media. Samples for *E. tarda* in TSB culture were collected at 0, 6, 18, and 24 h. (C) Screening of supernatants of *E. tarda* in TSB media. Abbreviations are as follows: IAld, indole-3-carboxaldehyde; IEt, tryptophol; IAM, indole-3-acetamide; IAA, indole-3-acetic acid; IAAld, indole-3-acetaldehyde; and IpyA, indole-3-pyruvate. Extracted ions were selected for IAld (m/z 145), IEt, (m/z 161), Indole (m/z 117), IAAld (m/z 159), IAM (m/z 174), IAA (m/z 175), and IpyA (m/z 203). (D) Chemical profiles of Trp-Indole derivatives from supernatants of various commensal bacteria in TSB medium for 1 day of cultivation. Values represent normalized production of Trp-Indole derivatives based on CFU. (E) Proposed model of *E. tarda* tryptophan catabolism. (F) EEC Gcamp fluorescence intensity in *Tg(neurod1:Gcamp6f)* zebrafish stimulated with different tryptophan catabolites. (G-H) Represented images (G) and quantification (H) of activated EECs in *Tg(neurod1:CaMPARI)* zebrafish that is stimulated with PBS or with CFS from *E. tarda* 23685 and *E. tarda* 15974. (I-J) Indole (I) and IAld (J) stimulation of Ca^2+^ influx in human TRPA1 expressing HEK-293T cells, measured as fluorescence increase of intracellular Calcium 6 indicator. (K-L) Effects of TRPA1 inhibition using various concentrations of inhibitor A967079, on subsequent Ca^2+^ influx in response to indole (100 µM, G) or, IAld (100 µM, H) in human TRPA1 expressing HEK-293T cells. Data are from a representative experiment performed in triplicate and repeated three times. (M-N) Sensitivity of mouse TRPA1 to indole and IAld. (M) Dose-response effects of indole and IAld (EC50 = 130.7 µM, 107.8 – 158.4 µM 95% CI for Indole; and, EC50 = 189.0 µM, 132.8 - 268.8 µM 95% CI for IAld). Concentration-response data were normalized to 1 mM cinnamaldehyde (CAD), a known TRPA1 agonist. (N) Effects of the Trpa1 inhibitor A967079, on [Ca^2+^]_i_ in response to 100 µM indole in mouse Trpa1-expressing HEK-293T cells. Cells were treated with A967079 before the addition of indole (100 µM). Changes in Calcium 6 fluorescence above baseline (Fmax-F0; maximal [Ca^2+^]_i_) are expressed as a function of Trpa1 inhibitor, A967079, concentration (IC50 = 315.5 nM, 202.3 – 702.3 nM 95% CI for indole). Concentration-response data were normalized to the response elicited by 100 µM Indole. Data represent mean ± s.e.m. of normalized measures pooled from two experiments, each performed in triplicate.

**Figure S8.**
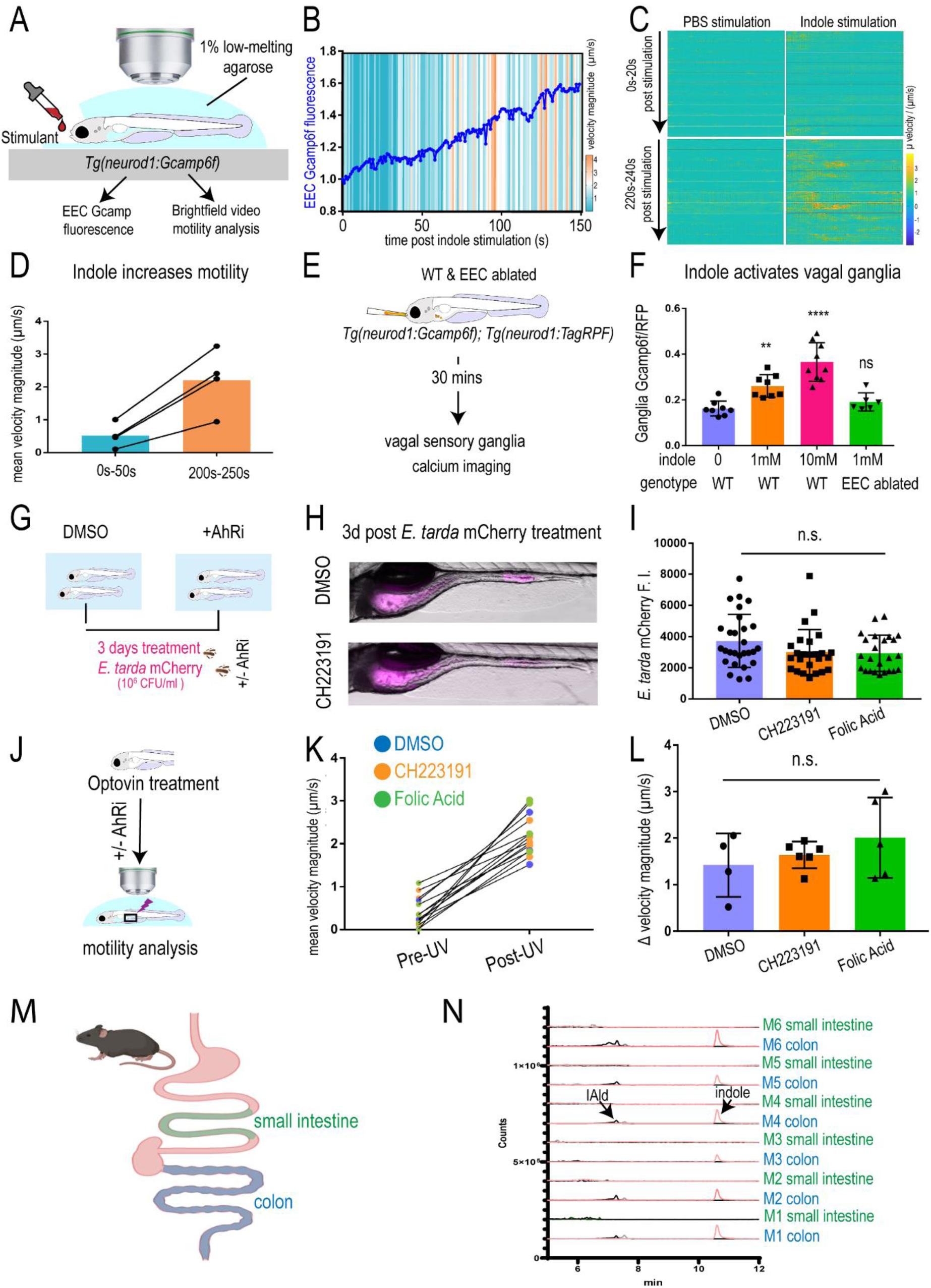
Effects of tryptophan catabolites and AhR inhibitor on intestinal motility, related to Main Figure 7. (A) Experimental model for measuring intestinal motility in response to indole stimulation. (B) EEC Gcamp6f fluorescence (blue line) and changes in intestinal motility (heat map) following indole stimulation. (C) Intestinal µ velocity in response to PBS or indole stimulation. (D) Mean intestinal velocity magnitude 0-50s and 200-250s following indole stimulation. (E) Schematic of experiment design in measuring the effects of indole or IAld in vagal ganglia calcium. WT or EEC ablated *Tg(neurod1:Gcamp6f); Tg(neurod1:TagRFP)* zebrafish that were gavaged with indole or IAld. (F) Quantification of the Gcamp6f to TagRFP fluorescence ratio in the whole vagal sensory ganglia in WT or EEC ablated zebrafish 30 minds following indole gavage. (G) Schematic of experiment design in testing the effects of AhR inhibitors on intestinal *E. tarda* accumulation. (H) Representative image of DMSO or AhR inhibitor CH223191 treated zebrafish that were infected with *E. tarda* expressing mCherry (*E. tarda* mCherry). (I) Quantification of *E. tarda* mCherry fluorescence in DMSO, AhR inhibitor CH223191 or Folic acid treated zebrafish intestine. (J) Schematic of experiment set up to examine the effects of AhR inhibitors in Trpa1^+^EEC induced intestinal motility. (K) Quantification of mean intestinal velocity magnitude in DMSO, CH223191 or Folic acid treated zebrafish before and post UV activation. (L) Quantification of mean intestinal velocity magnitude change in DMSO, CH223191 or Folic acid treated zebrafish up UV-induced Trpa1^+^EEC activation. (M) Schematic of small intestine and colonic regions in 10-week old SPF C57Bl/6 mice that were collected for HPLC-MS analysis. (N) Chemical profiles of Trp-Indole derivatives from colon and small intestine of conventionally-reared mice. Relative amounts of the Trp-metabolites from each mouse was normalized by tissue weight. M1-M3: males. M4-M5: females. Extracted ions were selected for Indole (m/z 117), IAld (m/z 145), and IEt, (m/z 161). One-way ANOVA with Tukey’s post test was used in F, I, L. ** P<0.01, **** P<0.0001, n.s. not significant.

