## Supplementary material for "Enteroendocrine cells sense bacterial tryptophan catabolites to activate enteric and vagal neuronal pathways": Key Resources Table

| REAGENT or RESOURCE | SOURCE | IDENTIFIER |
| --- | --- | --- |
| Antibodies | | |
| Chicken anti-GFP Polyclonal antibody | Aves Lab | Cat# GFP-1010, RRID:AB_2307313 |
| Living Colors® anti DsRed Polyclonal Antibody | TAKARA | Cat# 632496, RRID:AB_10013483 |
| Mouse anti-zebrafish gut secretory cell epitopes Monoclonal antibody [FIS 2F11/2] | Abcam | ab71286 |
| Mouse anti-zebrafish Zn-12 Monoclonal antibody | ZIRC | AB_10013761 |
| Rabbit anti-serotonin whole Polyclonal antibody | Sigma | Cat# 5545 |
| Mouse anti-p44/42 MAPK (Erk1/2) (L34F12) Monoclonal antibody | Cell Signaling | Cat# 4696 |
| Rabbit anti-Phospho-p44/42 MAPK (Erk1/2) (Thr202/Tyr204) (D13.14.4E) Monoclonal antibody | Cell Signaling | Cat# 4370T |
| Rabbit anti-chicken desmin Polyclonal Antibody | Sigma | Cat# D8281 |
| Rabbit anti-mouse PYY antibody | PMID: 28614796 | Custom antibody generated in Liddle Laboratory, aa4-21 (mouse) |
| Bacterial and Virus Strains | | |
| *Edwardsiella tarda* FL6-60 | PMID: 22003892 | N/A |
| *Edwardsiella tarda* LSE40; *pmkb:mCherry* | Gift by Mark Cronan | N/A |
| *Edwardsiella tarda* 23685 | ATCC | ATCC^23685^ |
| *Edwardsiella tarda* 15974 | ATCC | ATCC^15974^ |
| *Acinetobacter* sp. ZOR0008 | PMID: 26339860 | N/A |
| *Aeromonas veronii* ZOR0002 | PMID: 26339861 | N/A |
| *Shewanella* sp. ZOR0012 | PMID: 26339862 | N/A |
| *Enterobacter* sp. ZOR0014 | PMID: 26339863 | N/A |
| *Vibrio* sp. ZWU0020 | PMID: 26339864 | N/A |
| *Chryseobacterium* sp. ZOR0023 | PMID: 26339865 | N/A |
| *Exiguobacterium acetylicum* ZWU0009 | PMID: 26339866 | N/A |
| *Bacillus subtilis 168* | PMID: 18723616 | N/A |
| *Pseudomonas aeruginosa* PAK | PMID: 30524971 | N/A |
| *Plesiomonas* sp. ZOR0011 | PMID: 26339861 | N/A |
| *Escherichia coli* MG1655 | PMID: 26339862 | N/A |
| Chemicals, Peptides, and Recombinant Proteins | | |
| Optovin | Tocris Biotech | 4901 |
| HC030031 | Sigma | 377430-5g |
| Allyl isothiocyanate | Sigma | H4415-10MG |
| Atropine | Sigma | A0132 |
| 4-DAMP | Sigma | SML0255 |
| Clozapine | Sigma | C6305 |
| α-Bungarotoxin | Sigma | 203980 |
| Indole-3-carboxaldehyde | Sigma | 129445 |
| Indole-acetaldehyde | Ambeed | A626636 |
| Indole | Sigma | I3408-25G |
| L-Trytophan | Sigma | T8941-10mg |
| Indole-3-acetic acid sodium salt | Sigma | I5148-2G |
| Indole-3-pyruvic acid | Sigma | I7017 |
| Indole-3-acetamide | Sigma | 286281-1G |
| Tryptophol | Sigma | T90301-5G |
| CH030031 | Sigma | C8124-5M |
| Folic acid | Sigma | F7876-1G |
| Deposited Data | | |
| Raw and analyzed data | This paper | GSE151711 |
| Experimental Models: Cell Lines | | |
| HEK-293T cells | ATCC | ATCC-CRL-1573 |
| Experimental Models: Organisms/Strains | | |
| Zebrafish: *Tg(neurod1:CaMPARI)^rdu78^* | This paper | N/A |
| Zebrafish: *Tg(-5kbneurod1:Gcamp6f)^icm05^* | PMID: 27231612 | N/A |
| Zebrafish: *TgBAC(cldn15la:EGFP)^pd1034^* | PMID: 24504339 | N/A |
| Zebrafish: *Tg(-5kbneurod1:TagRFP)^w69^* | PMID: 22738203 | N/A |
| Zebrafish: *TgBAC(trpa1b:EGFP)^a129^* | PMID: 22190641 | N/A |
| Zebrafish: *Tg(neurod1:cre; cmlc2:EGFP)^rdu79^* | This paper | N/A |
| Zebrafish: *TgBAC(gata5:loxp-mcherry-stop-loxp-DTA)^pd315^* | This paper | N/A |
| Zebrafish: *trpa1b* mutant*^vu197^* | PMID: 18829968 | N/A |
| Zebrafish: *trpa1a* mutant*^hu2163^* | PMID: 18829968 | N/A |
| Zebrafish: *TgBAC(neurod1:EGFP)^nl1^* | PMID: 19424431 | N/A |
| Zebrafish: *Tg(-5kbneurod1:Gal4; cmlc2:EGFP)^rdu71^* | PMID: 31793875 | N/A |
| Zebrafish: *Tg(-5kbneurod1:Lifeact-EGFP)^rdu70^* | PMID: 31793875 | N/A |
| Zebrafish: *Tg(UAS:ChR2(H134R)-mCherry)^s1985^* | PMID: 26752076 | N/A |
| Zebrafish: *Tg(NBT:DsRed)^zf148^* | PMID: 29628374 | N/A |
| Zebrafish: *ret* mutant*^hu2846^* | PMID: 21490065 | N/A |
| Zebrafish: *sox10* mutant*^t3^* | PMID: 28207737 | N/A |
| Zebrafish: *TgBAC(chata:Gal4)^mpn202^* | PMID: 28701772 | N/A |
| Zebrafish: *Tg(UAS:Gcamp6s)^mpn101^* | PMID: 28701772 | N/A |
| Zebrafish: *Tg(UAS:NTR-mCherry)^c264^* | PMID: 24496182 | N/A |
| Zebrafish: *Tg(isl1:EGFP)^rw0^* | PMID: 23850871 | N/A |
| Zebrafish: *tph1b* mutant *^pd249^* | PMID: 28811310 | N/A |
| Zebrafish: *Tg(tph1b:mCherry-NTR)^pd275^* | This paper | N/A |
| Zebrafish: *Tg(β-act2:Brainbow1.0L)^pd50^* | PMID: 22538609 | N/A |
| Zebrafish: *Tg(neurod1:CaMPARI)^rdu78^* | This paper | N/A |
| Zebrafish: *Tg(-5kbneurod1:Gcamp6f)^icm05^* | PMID: 27231612 | N/A |
| Zebrafish: *TgBAC(cldn15la:EGFP)^pd1034^* | PMID: 24504339 | N/A |
| Zebrafish: *Tg(-5kbneurod1:TagRFP)^w69^* | PMID: 22738203 | N/A |
| Mouse: C57/BL6J | Jackson Laboratory | 000664 |
| Oligonucleotides | | |
| RT-qPCR for zebrafish gene: *pyyb*  NM_001327895 | Eurofins Genomics | AGCGTATCCACCCAAACCTG |
|  |  | GCCGGATGTCCTGTTCATCA |
| RT-qPCR for zebrafish gene: *ccka*  XM_001346104 | Eurofins Genomics | AACCAAAGGCTCATACCGCA |
|  |  | TCATATTCCTCGGCGCTTCG |
| RT-qPCR for zebrafish gene: *adcyap1a*  NM_152885 | Eurofins Genomics | GGGGTTTTCACGGACAGCTA |
|  |  | TGTGTCACAAAGCCGGGAAT |
| RT-qPCR for zebrafish gene: *insl5a*  NM_001037669 | Eurofins Genomics | TGCTGTAAGCAGACGAGACC |
|  |  | AGCAGAGGAACGTCAGGTCA |
| RT-qPCR for zebrafish gene: *fabp2*  NM_131431 | Eurofins Genomics | TGGAAAGTCGACCGCAATGA |
|  |  | TGAACTTGTCTCCGGTCTGC |
| RT-qPCR for zebrafish gene: *muc5.3*  XM_021477626 | Eurofins Genomics | ATGCGAACCATGGGGCTTTA |
|  |  | TTGTTCGCGTTCCCGTCATA |
| RT-qPCR for zebrafish gene: *sypa*  NM_001143977 | Eurofins Genomics | GATCGTGGCACCGTTTATGC |
|  |  | ATTGTAGCCTTGCTGGCTGT |
| RT-qPCR for zebrafish gene: *sypb*  NM_001030242 | Eurofins Genomics | ATCCTATGGGGAGGCAACCT |
|  |  | ACCTTCCTGTCCATAGCCCT |
| RT-qPCR for zebrafish gene: *agr2*  NM_001012481 | Eurofins Genomics | AGTGCTCTTGGTCATGGTGG |
|  |  | AGGGGCTTGTTCTTGGATCG |
| RT-PCR for zebrafish gene: *trpa1a*  NM_001007065 | Eurofins Genomics | TACCAACATGTCGTGTTTTCAGTG |
|  |  | GATTGCACACAACCGGTTTACA |
| RT-PCR for zebrafish gene: *trpa1b*  NM_001007066 | Eurofins Genomics | CTCATTTGTCTTGGAAAGGGAGC |
|  |  | GGAGGAAGTTGCGACCTGTT |
